# Joint experience-dependent representations of odors and temporal context in the zebrafish homolog of piriform cortex

**DOI:** 10.64898/2026.08.25.747023

**Authors:** Tommaso C. Caudullo, Jaap van Krugten, Jan Eckhardt, Rainer W. Friedrich

## Abstract

Intelligent behavior requires experience-dependent internal representations of relevant information. We examined representational learning in telencephalic area pDp of adult zebrafish, the homolog of piriform cortex. Activity was measured by multiphoton imaging through a prism in head-fixed fish trained in an odor discrimination task. Population dynamics could be decomposed into independent modes, each consisting of distributed activity across mixed-selectivity neurons, that encoded odor identity, novelty, and the progression of time. Training sharpened novelty detection, strengthened odor-specific response components, and geometrically untangled representational manifolds. Experience-driven plasticity generalized across chemically related stimuli. Main observations could be reproduced phenomenologically by a model assuming linear interactions between novelty-, identity- and time-encoding activity modes. We conclude that pDp jointly represents olfactory and temporal information relevant for episodic memory. Experience enhances the accessibility and generalization of information, indicating that pDp supports brain-wide learning by geometrically organizing olfactory and time-related information on neural manifolds.

## 1 Introduction

Interpreting the stream of sensory input is essential for intelligent behavior and requires the flexible, experience-dependent classification of high-dimensional activity patterns. These brain functions are believed to depend on autoassociative memory networks that store information about relevant structure in the world in their connectivity matrix. Autoassociative memory has been proposed to be a main function of piriform cortex (PCx) and its teleost homologue, telencephalic area pDp (posterior part of the posterior zone of the dorsal telencephalon) (Haberly and Bower, 1989; Li and Hertz, 2000; Haberly, 2001; Mueller et al., 2011; Wilson and Sullivan, 2011; Miyasaka et al., 2014; Rupprecht and Friedrich, 2018).

PCx/pDp are major targets of the olfactory bulb, where primary odor information is pre-processed and represented by distributed activity patterns (Haberly and Price, 1977; Friedrich and Laurent, 2001; Wiechert et al., 2010; Miyasaka et al., 2014; Chae et al., 2022; Hernandez et al., 2025). Moreover, PCx/pDp interact with other cortical and subcortical brain areas including lateral entorhinal cortex and prefrontal areas (Haberly, 2001). The distributed organization of afferent and recurrent (“associational”) connectivity (Yaksi et al., 2009; Franks et al., 2011; Ghosh et al., 2011; Miyamichi et al., 2011; Sosulski et al., 2011; Miyasaka et al., 2014; Fink et al., 2025) gave rise to the hypothesis that neurons in PCx/pDp integrate diverse, possibly arbitrary, combinations of local and long-range inputs. Recurrent connections exhibit activity-dependent plasticity and metaplasticity (Hasselmo and Barkai, 1995) and account for a substantial fraction of the total synaptic input to PCx/pDp neurons (Poo and Isaacson, 2009, 2011; Rupprecht and Friedrich, 2018). Odors evoke widespread activity in PCx/pDp without obvious chemotopic organization (Illig and Haberly, 2003; Rennaker et al., 2007; Stettler and Axel, 2009; Yaksi et al., 2009; Blumhagen et al., 2011; Roland et al., 2017; Pashkovski et al., 2020) . These observations are consistent with the hypothesis that PCx/pDp function as autoassociative networks that establish representations of arbitrary odor objects by experience-dependent modifications of recurrent synaptic connectivity between chemical feature detectors (Haberly and Bower, 1989; Li and Hertz, 2000; Haberly, 2001; Wilson and Sullivan, 2011; Babadi and Sompolinsky, 2014).

This classical hypothesis predicts that input patterns related to learned odor objects are mapped onto common output patterns by pattern completion (Wilson and Sullivan, 2011). In autoassociative memory models, pattern completion can be a consequence of convergent dynamics towards attractor states that are defined by experience-dependent recurrent connectivity. Although experimental approaches showed that odor exposure and associative learning modify odor responses in PCx/pDp (Schoenbaum and Eichenbaum, 1995; Kadohisa and Wilson, 2006; Chapuis and Wilson, 2012; Shakhawat et al., 2014; Pashkovski et al., 2020; Schoonover et al., 2021; Poo et al., 2022; Fink et al., 2025; Hu et al., 2026), large-scale population activity measurements failed to provide obvious evidence for distinct attractor states (Roland et al., 2017; Pashkovski et al., 2020; Schoonover et al., 2021; Fink et al., 2025; Hu et al., 2026). Nonetheless, odor sampling or olfactory discrimination training enhanced the ability to decode the identity of familiar and related stimuli from odor-evoked population activity (Pashkovski et al., 2020; Poo et al., 2022; Fink et al., 2025; Hu et al., 2026). These observations suggest that experience reorganizes the internal representation of odor space by geometrical modifications of activity subspaces (“neural manifolds”) without establishing discrete attractor states (Meissner-Bernard et al., 2025b, 2025a).

Odor representations in PCx/pDp are modified by experience on multiple timescales. Prolonged or repeated odor stimulation decreases response amplitude and changes activity patterns by mechanisms that are not accounted for by sensory adaptation in the nose or olfactory bulb (Wilson, 1998, 2000; Kadohisa and Wilson, 2006; Jacobson et al., 2018). In pDp of adult zebrafish, for example, repetitive brief odor stimuli separated by 2 min resulted in an NMDA receptor-dependent attenuation and reorganization of odor responses that recovered within tens of minutes (Jacobson et al., 2018). pDp therefore holds odor information in a short-term memory buffer that enables computations such as novelty detection. In anterior PCx of mice, odor representations drift on timescales of days and weeks (Schoonover et al., 2021), similar to representational drift of spatial cognitive maps in the hippocampus (Rubin et al., 2015). This drift was slowed down by odor exposure, suggesting that PCx maintains non-stationary representations of odor space that are updated by slow changes in the olfactory environment (Schoonover et al., 2021). Additional observations show that odor representations in PCx/pDp and related areas undergo specific and presumably long-lasting changes after odor discrimination training (Chapuis and Wilson, 2012; Shakhawat et al., 2014; Meissner-Bernard et al., 2019; Hernandez et al., 2025; Hu et al., 2026; Schiltz et al., 2026). These observations raise the question how PCx/pDp integrate information about odors and time in an experience-dependent manner.

To address these questions we trained adult zebrafish in an odor discrimination task and measured responses to repeated applications of multiple conditioned and novel odors. A minimally invasive method using an optical prism was developed to measure activity in pDp of awake head-fixed fish by two-photon calcium imaging. We found that information about novelty and identity of odors was encoded by distinct components of activity patterns across populations of neurons with mixed selectivity. No reward-related changes in odor representations were observed after training. However, training accelerated the initial dynamics of odor responses, increased response reliability, enhanced the odor-specificity of the novelty-related response component, and resulted in an untangling of representational manifolds (Chou et al., 2025). These effects generalized from representations of conditioned odors to related novel odors. We further identified an odor-independent component of population activity that is present in the absence of stimulation and provides information about the progression of global or episodic time. These results reveal that multiple stimulus variables, notably novelty, identity and relative time, are jointly represented in pDp by different modes of neuronal population dynamics. Information about task-relevant odors is not stored by discrete object representations but in modifications of the internal map of odor space, enhancing the decodability of task-relevant information and facilitating future learning.

## 2 Results

### 2.1 Odor-evoked activity in pDp of adult zebrafish in vivo

Previous measurements of odor-evoked activity in pDp (Yaksi et al., 2009; Blumhagen et al., 2011; Jacobson et al., 2018; Rupprecht and Friedrich, 2018; Hu et al., 2026) were performed in an ex-vivo preparation of the juvenile or adult zebrafish brain (Zhu et al., 2012). In vivo, activity has been measured in dorsal telencephalic areas of head-fixed adult zebrafish by 2-photon imaging through the intact skull (Huang et al., 2020; Torigoe et al., 2021; Palacios-Flores et al., 2026) but optical access to pDp is complicated by its location in the ventro-lateral pallium, typically *>*500 mm below the dorsal surface. We therefore replaced a bone between the brain and the eyecup by a prism to obtain lateral optical access to the telencephalon (Fig. 1a and Fig. S1a-b,d). Using a transgenic line expressing GCaMP6f throughout the adult telencephalon (Tg[neurod1:GCaMP6f]; Rupprecht et al., 2016, Fig. S1c), this approach allowed us to measure activity across 97 – 301 pDp neurons (mean ± SD: 215 ± 41 neurons; N = 35 fish) in sagittal planes by 2-photon calcium imaging in awake behaving fish (Fig. 1a-c). Somatic calcium signals (ΔF/F) were transformed into firing rate estimates using CASCADE (Rupprecht et al., 2021, Fig. 1c and Fig. S1e).

**Figure 1.**
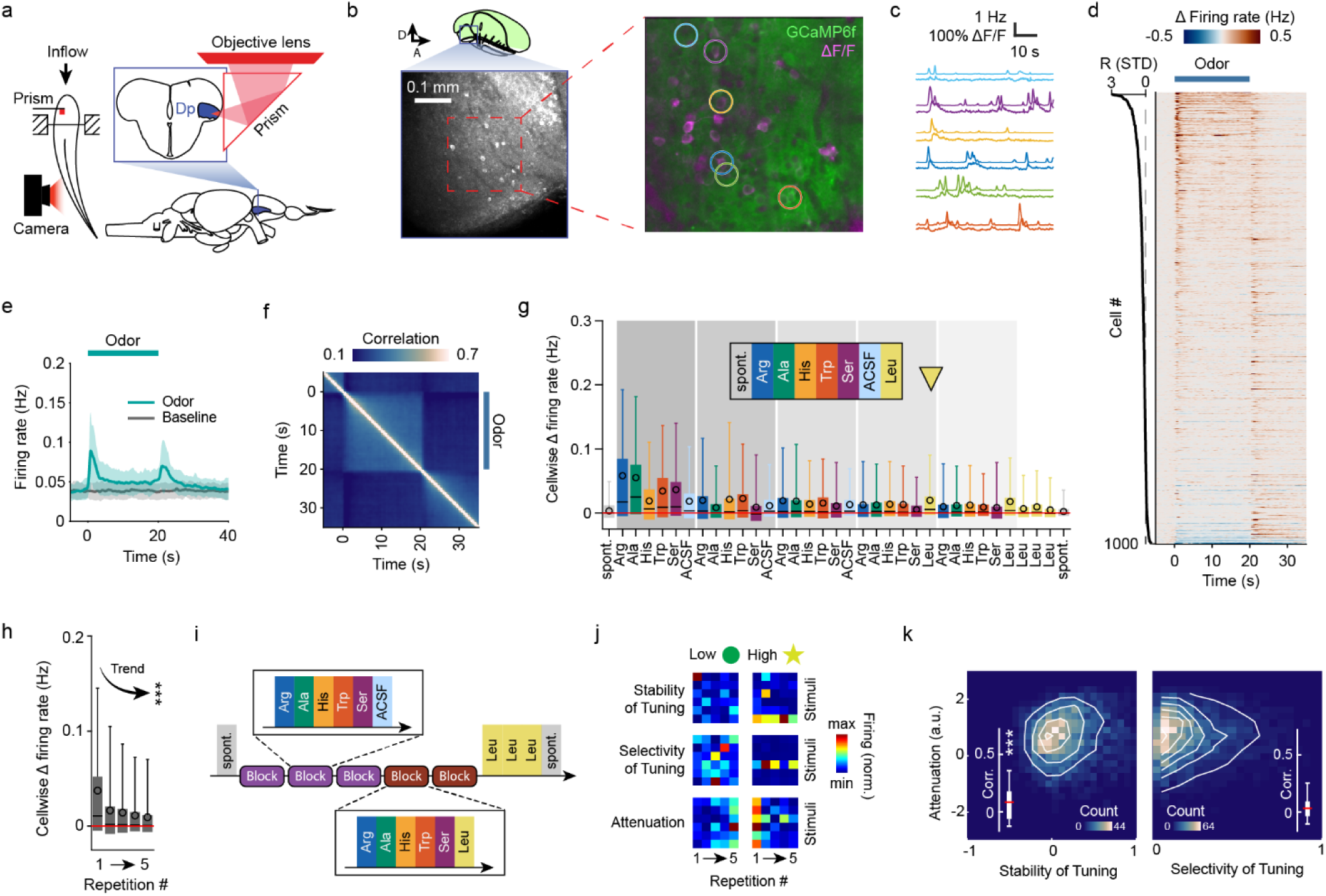
Responses to odors in adult pDp of Naïve fish. **a** Left: Overview of experimental setup and microprism implantation. **b** View of pDp through the prism. Top left: sagittal diagram of forebrain anatomy, adapted from (Wullimann et al., 1996), at a depth of 50 μm. Bottom left: maximum intensity projection of one representative low-zoom anatomical stack (see also Fig. S1). Right: time-averaged raw fluorescence (green) and response to an odor (ΔF/F; magenta). **c** ΔF/F (bottom traces) and inferred firing rate (top traces) of the cells circled in **b**. **d** Trial-averaged baseline-subtracted responses of 1000 randomly selected pDp neurons to all odors from Naïve fish, sorted by their odor responsiveness score (R, left). Baseline was taken as the interval from 5s to 1s before odor onset. Bar indicates odor presentation. **e** Average single-cell peri-stimulus time trace across Naïve fish (mean + SD over n = 14 single-fish trial-averaged traces). **f** Pearson correlations of single-trial population activity across time around the odor pulse, averaged over all odor responses and fish (n = 1050 responses, see Fig. S10i for Naïve only). **g** Mean firing rate change between a 20 s pre-stimulus window and the odor response in consecutive trials (n = 2892 cells, pooled data from all Naïve fish). Mean responses were positive but ∼25% of cells showed negative responses in each trial. Note stronger responses to initial odor stimulations and a slightly increased response to the first application of Leu (yellow arrow, see Fig. S4d-e for quantification). Shaded gray backgrounds show trial blocks 1-5 (see **i**). **h** Average single-stimulus firing rate change across repetitions, across all odors in Naïve fish. Friedman test for significant trends: p < 2.2·10^-308^. **i** Trial sequence. Odor trials were organized in blocks of 6 trials. Leu is presented first in block 4 and repeated 3 more times after block 5. Each amino acid was thus presented 5 times. **j** Six example cells, scoring low (top row) or high (bottom row) for attenuation, selectivity of tuning and stability of tuning. Each subplot indicates the cell’s average firing rate during the response to the corresponding stimulus and repetition. **k** Relationship between single-cell attenuation and tuning metrics. White lines show smoothed isoclines. A linear mixed-effects model was applied to correct for small subject-specific biases. Spearman correlation of attenuation vs stability of tuning: r = 0.077, p = 1·10^-4^. Attenuation vs selectivity of tuning: r = 0.032, p = 0.089. ***: p-value *<*0.001. Box plots: 25^th^ and 75^th^ percentile; whiskers: mean + SD; horizontal line: median; black circle: mean.

Head-fixed fish were maintained in the dark and odors were applied for 20 s through a constant flow of buffered medium. Odors evoked responses in stimulus-specific subsets of neurons that typically followed a phasic-tonic time course with an off-response (Fig. 1d-e). The Pearson correlation between activity patterns evoked by the same odors at different times or in repeated applications was modest (Fig. 1f and below), indicating considerable variability. No obvious topographic organization of odor responses was detected. These observations are consistent with previous ex-vivo measurements of odor-evoked activity in pDp (Blumhagen et al., 2011; Jacobson et al., 2018; Hu et al., 2026). The most obvious difference between ex vivo and in vivo activity was a higher baseline activity in the in vivo condition (see below).

To further characterize basic features of odor-evoked activity we quantified the responsiveness of individual neurons as the difference between firing rates during odor presentation and the mean rate over the entire recording session, normalized by the standard deviation of rates averaged over each trial ([response – session_mean]/SD[trials]). Responsiveness was biased towards positive values, indicating that most firing rates increased during odor stimulation (Fig. S3a). Some neurons showed activity modulations related to tail motion but this activity was nearly uncorrelated to odor-evoked activity (Fig. S3). As observed previously, odor-evoked increases in firing rates were most prominent during a transient initial response phase that was followed by lower, more sustained mean activity (Fig. 1e). Correlations between activity patterns measured at different timepoints decayed rapidly on short timescales but remained positive throughout stimulus presentation (Fig. 1f). Substantial positive correlations were also observed between population activity patterns before and after the odor stimulus, with sharp transitions at stimulus onset and offset (Fig. 1f, Fig. S10).

Odors were applied in five blocks of six different stimuli (amino acids [10^-4^ M] or a blank control [ACSF]). As observed ex-vivo (Jacobson et al., 2018), odor responses decreased with repeated stimulation (inter-trial interval: 3 min). This attenuation was present within the first block but most pronounced between the first and the second block (Fig. 1g; Fig. S4a). These observations indicate that response attenuation was odor-selective, with some cross-attenuation between odors, and most pronounced between the first and the second repetition of an odor (Fig. 1g-h, Fig. S4a).

To further assess the odor-specificity of this attenuation we changed the sequence of odor stimuli after block 3 (Fig. 1i): in the first three blocks, five amino acids (Arg, Ala, His, Trp, Ser) and one blank control (ACSF) were applied in the same sequence while in blocks 4 and 5, ACSF was replaced by a novel amino acid (Leu). Block five was then followed by three further repetitions of Leu. This protocol allowed us to compare responses to different odors (within blocks) and responses to repeated presentations of the same odors (across blocks). Moreover, responses to novel odors could be compared after prolonged absence of odor stimulation (first block) and after prior stimulation with other odors (first Leu application in block 4). Furthermore, repeated odor responses could be compared with intervening presentations of other odors (across blocks) or without (repeated applications of Leu).

Comparing responses across blocks showed that the amplitude and variance of the population response decreased significantly, particularly after the first presentation of each odor (Fig. 1h, S4a, S8c). Odor responses therefore contain a component that is specific for novel stimuli and attenuates over repetitions. To quantify response attenuation of individual neurons we defined an “attenuation score” that measures the relative reduction in odor-evoked firing across repetitions (Fig. 1j, Fig. S4c; see section 4). Attenuation varied substantially across cells (Fig. S4c), and was only weakly correlated to their average firing rate (Fig. S4b-c), indicating that attenuation cannot be explained by global adaptation. We further asked whether attenuation is systematically related to other features of odor responses, namely the selectivity and stability of odor tuning (Fig. 1j, Fig. S4c, Methods). Tuning selectivity of individual neurons was measured by lifetime sparseness while tuning stability was quantified by the mean correlation between tuning curves across repetitions (see Methods). Attenuation scores were not correlated to tuning selectivity (Fig. 1k, Fig. S4c) and only weakly correlated with tuning stability (r = 0.077, p = 1·10^-4^)(Fig. 1k, Fig. S4c), indicating that attenuation is largely unrelated to tuning. These results suggest the attenuation of odor responses is represented by a distinct mode of population activity that represents novelty (see below).

### 2.2 Experience-driven changes in odor representations: basic observations

We next trained adult zebrafish in an established odor discrimination task (Namekawa et al., 2018; Hu et al., 2026) (Fig. 2a). Briefly, individual fish were exposed to each of two odor stimuli (CS+, CS-) nine times per day for 30 s. CS+ but not CS-was followed by food delivery at a specific location. Appetitive behavior was quantified during odor presentation by an integrated measure combining multiple behavioral components (Namekawa et al., 2018, and Fig. S5a-b). After 5 days of training, appetitive behavior was significantly more pronounced during presentation of the CS+ than the CS-(Fig. 2c), as observed previously (Namekawa et al., 2018; Frank et al., 2019; Hu et al., 2026).

**Figure 2.**
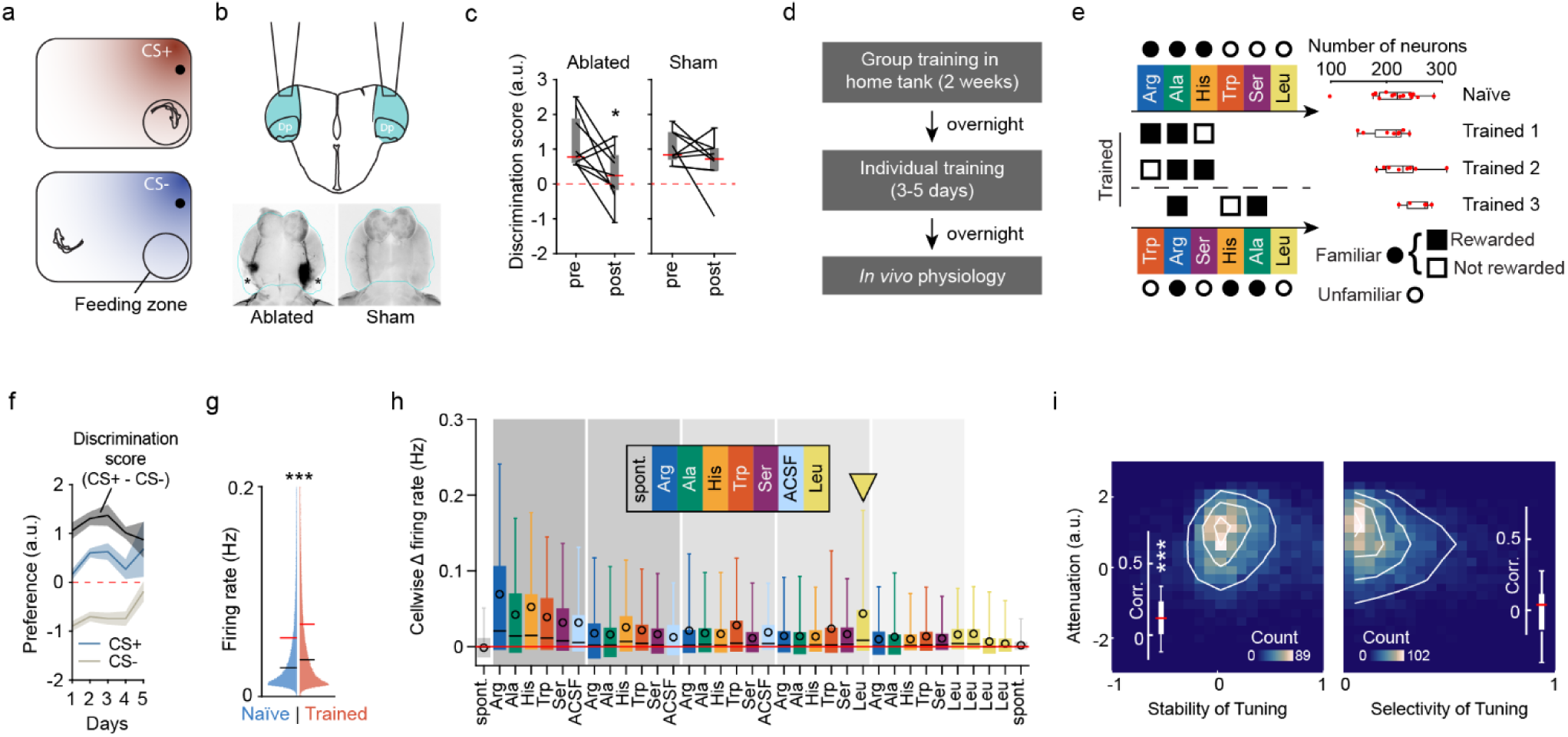
Effects of odor discrimination training on responses of individual neurons. **a** Individual training, schematic: experienced animals respond appetitively during the presentation of the CS+, but not the CS-. **b** Top: schematic: bilateral ablation of Dp and overlying telencephalic tissue. Bottom: maximum intensity projection from a 3D confocal stack of the cubic-cleared forebrain. Left: ablated brain. Outlines depict original shape to show ablated regions. Black regions indicate a high density of extant damaged tissue. In the example, this includes a region of the dorsolateral subpallium, known to project to pDp and a small neighborhood of pDp with fiber tracts to the OB. Right: example forebrain from a sham surgery animal (craniotomy without aspiration of Dp). **c** Effect of bilateral Dp ablation on behavioral preference. Red dotted line: no average preference. After ablation, behavioral preference for the CS+ was significantly decreased. Right: sham surgery had no statistically significant effect on discrimination. Paired 1-sided t-test, ablated p = 0.016, sham p = 0.097. **d** Schematic: training schedule for physiological recordings. **e** Left: schematic of odor-reward associations and trial block structure for each experimental group. Right: number of recorded neurons per fish for each group. **f** Combined preference scores for the CS+ and CS- in experienced fish, including the combined discrimination score (CS+ - CS-), over days (individual training/testing). All fish were pre-trained as a group in the home tank. Lines: mean ± s.e.m. **g** Mean firing rates during odor presentation pooled across all trials and cells, clipped at 0.2 Hz. Horizontal black lines show median (difference of medians: 0.008 Hz); red lines show mean (difference of means: 0.013 Hz). 2-sided 2-sample KS test: p < 2.2·10^-308^). **h-i** Same as in Fig. 1 for trained fish (see also Fig. S4). **h** n = 4635 cells. **i** Spearman correlation of attenuation vs stability of tuning: r = 0.080, p = 8.64·10^-8^. Attenuation vs selectivity of tuning: r = -0.011, p = 0.463.

To explore whether discrimination learning depends on pDp we lesioned trained fish by aspiration through a micropipette. This procedure completely removed pDp but also affected adjacent dorsal telencephalic areas (Fig. 2b). Odor discrimination tested on the following day was abolished in lesioned but not in sham-operated fish (Fig. 2c), supporting the hypothesis that pDp is involved in the learned discrimination between similar odors.

To examine how training modifies odor-evoked activity we made two modifications to the behavioral task (Fig. 2d-f). First, prior to discrimination training of individual fish and quantification of appetitive behavior, fish were pre-trained as groups in their home tank for two weeks (Hu et al., 2026). Second, fish were trained to discriminate two rewarded odors (CS+1, CS+2) from one non-rewarded odor (CS-; five CS+1, four CS+2 and nine CS-trials per day). In different training groups (“Trained 1”, “Trained 2”, “Trained 3”), we exposed fish to the same three odors (Arg, Ala, His) but varied their reward assignments (CS+1, CS+2, CS-; Fig. 2e) to assess whether experience-dependent changes in odor representations depend on behavioral outcome (Figs. S6, S7 and S9f-h). Appetitive behavior was significantly more pronounced in response to CS+1 and CS+2 than CS- in all training groups, demonstrating that fish learned the task (Fig. 2f, Fig. S5a-b).

After training we measured responses to the same six amino acid odors as in Naïve fish, thus comprising three familiar (Arg, Ala, His) and three unfamiliar stimuli (Trp, Ser, Leu). The sequence of odor stimuli within blocks was the same in all groups except for “Trained 3”: this group received the same behavioral training as “Trained 1” but stimulus blocks started with an unfamiliar odor (Trp) to assess potential effects of stimulus sequence (Fig. 2e). However, as differences between Trained groups 1 – 3 were minor, results were pooled for further analyses unless noted otherwise.

The distribution of mean firing rates, averaged over the duration of odor presentation, was skewed and shifted significantly towards higher frequencies in Trained fish (Fig. 2g, Fig. S5c). Differences in firing rates between Naïve and Trained fish were particularly prominent during the initial phases of on- and off-responses (Fig. 1e, Fig. S5d). As in Naïve fish, response amplitudes declined over trials and responses to the first presentation of each odor were significantly larger than subsequent responses (Fig. 2h, Fig. S5e). Firing rate changes of individual neurons were negatively correlated to the magnitude of odor responses on the first repetition but this correlation was very low. No significant correlation was observed on subsequent repetitions (Fig. S4f-g). As in Naïve fish, attenuation was not correlated with tuning stability but weakly correlated with tuning selectivity (Fig. 2i, Fig. S4h). Trained fish therefore exhibited a prominent novelty-related attenuation of odor responses, consistent with Naïve fish.

### 2.3 Experience enhanced the specificity of novelty detection

To further analyze the representation of novelty we surmised that novelty-related activity should fulfill the following requirements: (1) the response to the first repetition of each odor should be larger than responses to subsequent repetitions, and (2) the first response to a novel odor should exceed responses to other odors that had been repeated before. These predictions were tested by analyzing responses to Leu, the stimulus that was introduced only at the end of block 4 (Fig. 1i). Consistent with these predictions, the first response to Leu was significantly larger than subsequent responses to Leu, and significantly larger than responses to other stimuli that had been applied previously (Fig. 3a, Fig. S4d,e,i,j). The component of the population activity undergoing attenuation does therefore represent odor-specific novelty, at least in part. We further observed that the first response to Leu and the subsequent attenuation were more pronounced in Trained fish (Fig. 3a-b, Fig. S4e,j). Training therefore enhanced novelty-related activity following previous stimuli, indicating a higher odor-specificity of novelty detection.

**Figure 3.**
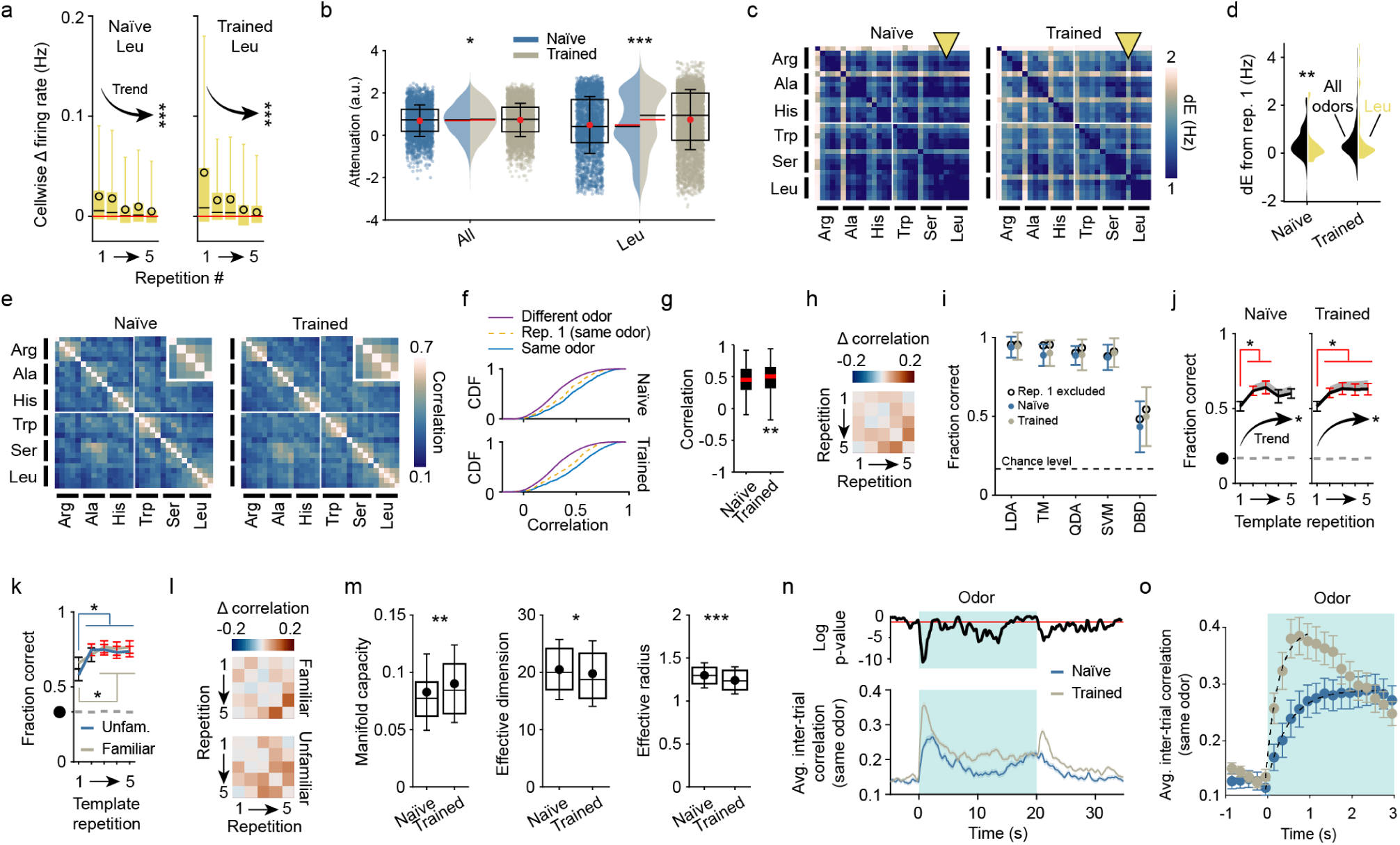
Effects of training on population activity. **a** Average firing rate change during repeated responses to Leu across training conditions. Friedman test for trends: p(Naïve) = 5.86·10^-80^; p(Trained) = 3.56·10^-141^. **b** Single-cell attenuation scores, computed based on all odorants (left) or only Leu (right). Attenuation is slightly higher in Trained fish. This difference is prominent for Leu. Pairwise Mann-Whitney U test: ‘All’ p = 0.043; ‘Leu’ p = 5.02·10^-17^. **c** Euclidean distance (dE) across single- trial time-averaged response vectors in Naïve and Trained fish. Matrices are sorted by stimulus identity. Yellow arrowhead: first Leu trial. **d** Odor-specificity of the pairwise dE between each first repetition of an odor and every other repetition. Black: all odors combined; yellow: Leu only. While dE from trial 1 is as high for Leu as for other odors in Trained fish (Pairwise Mann-Whitney U test, p = 0.055), it is significantly lower for Leu in Naïve fish (p = 1.6·10^-3^). This indicates a higher odor specificity of the novelty effect in Trained fish. **e** Pearson correlation across single-trial time-averaged response vectors in Naïve and Trained fish. Inset: correlations across same-odor repetitions, averaged over stimuli. **f** Cumulative distribution of correlation coefficients between trials of the same odor (solid blue), different odors (solid purple) or between repetition 1 and other repetitions of the same odor (dashed yellow). Top: Naïve; bottom: Trained. Mann-Whitney U test, with FDR correction: same vs different, p(Naïve) = 1.64·10^-58^, p(Trained) = 9.22·10^-108^; same vs rep 1, p(Naïve) = 3.00·10^-5^, p(Trained) = 3.67·10^-10^; different vs rep 1, p(Naïve) = 4.64·10^-10^, p(Trained) = 4.36·10^-18^. **g** Correlations between trials of the same odor in Naïve and Trained fish. Mann-Whitney U test, p = 6.5·10^-3^. **h** Difference between mean inter-trial correlation matrices between same-odor trials in Trained and Naïve fish (insets in **e**). Most correlations were slightly higher in Trained fish, particularly for late repetitions. **i** Odor identity classification accuracy, based on the response-averaged single-trial activity vectors using different linear classifiers, each trained on all trials. LDA: linear discriminant analysis; TM: template matching (to repetition-averaged template patterns, using Pearson correlation as a distance metric); QDA: quadratic discriminant analysis; SVM: support vector machines; DBD: direct basis decoders (Methods). **j** Fraction of correct template matching classifications in Naïve (left) and Trained (right) fish, using single trial response patterns as templates (x-axis) and testing on all remaining repetitions of each odor (59±14% for Naïve, 60±17% for Trained; n = 14 Naïve fish or 21 Trained fish; paired Wilcoxon signed rank tests between performance using repetition 1 or another repetition as template. Red error bars indicate p*<*0.05). Black circle: chance level; gray dashes: average performance of 50 equivalent decoders using shuffled cell IDs; gray shaded area: magnitude of performance increase when excluding repetition 1 trials. Trends: Friedman test. See also Fig. S9c. **k** Template matching performance in Trained fish, using only familiar or unfamiliar odors. **l** Difference between inter-trial correlation matrices for the same odor in Trained and Naïve fish, focusing on Arg, Ala and His (familiar in Trained fish) or on Trp, Ser, Leu (unfamiliar). Inter-trial correlations are higher in Trained fish for both familiar and unfamiliar odors. **m** Quantification of odor manifold capacity and geometrical features. Mann-Whitney U test: Capacity, p = 4.1·10^-3^; Dimension, p = 0.021; Radius, p = 3.00·10^-5^. See also Fig. S11. Box plots and whiskers: quartiles and standard deviations; horizontal lines: median; black circle: mean. **n** Bottom: same-odor inter-trial correlations on a sliding time window in Naïve and Trained fish (mean ± s.e.m.), averaged over all odors. Top: Log p-values of same-bin comparison across groups (Mann-Whitney U test). Red line: significance level (p*<*0.05). **o** Onset dynamics of inter-trial correlations between trials of the same odor (mean ± s.e.m.). Dashed line: exponential fit on the ascending phase.

To further analyze the specificity of novelty detection we examined Euclidean distances between odor response patterns across all trials. Each response pattern was represented by an activity vector representing firing rates of individual pDp neurons, averaged over the stimulus duration. Trials were re- ordered by odor identity to compare successive responses within and across odors. Overall, Euclidean distances were higher in Trained fish, consistent with the modest increase in firing rates (Fig. 3c). Moreover, in Trained fish, the responses to the first presentation of each odor were more different from other responses (higher Euclidean distance, Fig. 3d). In Naïve fish, a high distance between the first and subsequent responses to the same and other odors was seen only for the first two stimuli (Arg and Ala). First-trial responses to other odors of block 1 (His, Trp, Ser) were only slightly more distant from subsequent responses, and no obvious difference was observed between the first and subsequent responses to Leu (Fig. 3c-d; yellow arrowhead depicts first response to Leu). Hence, prior odor stimulation reduced novelty-related responses to other odors, implying cross-talk between novelty-related response components. In Trained fish, in contrast, the first trial-response to all odors including Leu was substantially different from subsequent responses to the same odor and responses to other odors (Fig. 3c-d). Hence, prior odor stimulation had little effect on novelty-related responses, further demonstrating that the novelty component was more selective in Trained fish.

### 2.4 Experience enhanced decodability of odor identity

To further analyze experience-driven changes in odor representations we quantified the similarity between activity vectors, averaged over the duration of stimulus presentation, by the Pearson correlation (Fig. 3e). Responses to repeated presentations of the same odor were higher than correlations between responses to different odors but trial-to-trial correlations rarely reached high values (*>*0.6; Fig. 3e,f), consistent with previous observations in PCx/pDp (Roland et al., 2017; Frank et al., 2019; Pashkovski et al., 2020; Hu et al., 2026). Correlations between same-odor trials were slightly higher in Trained fish (Fig. 3g,h), indicating that training slightly reduced trial-to-trial variability of time-averaged odor responses. To analyze effects of experience on pattern correlations in more detail we averaged correlations between the five repetitions across odors (Fig. 3e, insets). Correlations between the first and subsequent trials were consistently lower than subsequent correlations, confirming that odor representations were not only scaled but also reorganized after the first response. Furthermore, correlations between subsequent responses increased over trials, particularly in Trained fish. Correlations between responses to different odors, in contrast, did not change significantly across repetitions (Fig. S9a). Hence, experience increased the reliability of odor responses on a timescale of minutes, particularly in Trained fish (Fig. 3h).

In Trained fish, correlations between responses to familiar odors (Arg, Ala, His) were indistinguishable in training groups with different reward assignments (Fig. S6). Moreover, trial-to-trial correlations were not obviously different between responses to the two rewarded odors (CS+1, CS+2) or responses to rewarded and non-rewarded (CS-) odors (Fig. S9b). Hence, no systematic effect of task outcome (reward) was detected on the similarity between odor representations.

Representations of odor identity were further analyzed by training different classifiers (linear discriminant analysis [LDA], template matching [TM], quadratic discriminant analysis [QDA], support vector machine [SVM], direct basis decoder [DBD]; see Methods) to decode odor identity from activity vectors pooled over all trials (Fig. 3i). Using all classifiers, decoding performance was slightly higher in Trained fish, both for familiar and unfamiliar odors, but this trend was not statistically different (Fig. 3i). Because identity decoding may be complicated by novelty-related response components during the initial odor presentations we repeated the same analyses after excluding the first repetition of each odor. As expected, decoding performance was slightly increased but subtle differences between Naïve and Trained fish were preserved (Fig. 3i).

We further decoded stimulus identity by template matching using odor responses in successive repetitions as templates (Fig. 3j). Mean decoding performance was lowest based on response patterns evoked by the first odor application, consistent with the contribution of a prominent novelty component. In subsequent repetitions, decoding performance increased significantly both in Naïve (Friedman test: p = 0.027, see also individual inter-repetition comparisons) and in Trained fish (Friedman test: p = 0.012, see also individual inter-repetition comparisons). Similar observations were made using a support vector machine (Fig. S9c). Furthermore, decoding became more accurate when later repetitions were used as targets, independent of whether templates were generated from early or late repetitions (Fig. S9d). These observations were more consistent and pronounced in Trained fish (Fig. 3j; Fig. S9c,d). Hence, information about odor identity could be retrieved more reliably after priming of pDp by previous odor exposure, particularly in Trained fish. This observation can be explained, at least in part, by the attenuation of the novelty-related response component.

In Trained fish, distances and correlations were similar between activity patterns evoked by familiar and unfamiliar odors (Fig. 3c,e). Consistent with this observation, no difference in decodability of familiar and unfamiliar odors was observed (Fig. 3k) and correlations between activity patterns evoked by the same odors increased similarly over repetitions (Fig. 3l). Hence, effects of training generalized across representations of related odors.

As commonly used classifiers are limited in their ability to decode information from geometrically complex neural manifolds (DiCarlo and Cox, 2007; Chung and Abbott, 2021) we further analyzed decodability using “Geometry Linked to Untangling Efficiency” (GLUE) theory. This mathematical framework quantifies untangledness of potentially high-dimensional manifolds—i.e., their separability—by the measure of manifold capacity (Chou et al., 2025). This measure represents the amount of linearly decodable information per neuron and is directly linked to manifold geometry and neuronal covariability, thus providing direct insights into features of neural manifolds relevant for classification (Fig. S11a). At the same time, manifold capacity has a direct computational interpretation because it is mathematically linked to the efficiency, capacity and robustness of pattern classification (Chung et al., 2018; Chou et al., 2025).

A neural manifold representing an odor was defined as the distribution of activity vectors (“point cloud”) measured in individual time points (image frames) during odor presentation, pooled over all time points and repetitions (unless noted otherwise). Hence, neural manifolds were not constructed from time-averaged activity but each point in the point cloud represented the population activity measured at a given time point in a given trial. As a consequence, neural manifolds could, in principle, be high-dimensional and geometrically complex (Fig. S11a). Ex-vivo activity measurements revealed that manifold capacity of odor responses in pDp was increased after training in a related two-odor discrimination task (Hu et al., 2026). Consistent with these results, we found that manifold capacity measured in vivo was significantly higher after training in our 3-odor discrimination task. Experience therefore resulted in an untangling of representational manifolds, thus enhancing the accessibility of information relevant for odor identity decoding.

Standard GLUE theory expresses manifold capacity as a function of four interpretable parameters that represent two geometrical properties of manifolds and their correlations (Chou et al., 2025): (1) the effective manifold radius (the “compactness” of a manifold), (2) the effective manifold dimension (the “flatness” of a manifold), (3) the effective center alignment (the “pattern correlation” between activity vectors representing manifold centers), and (4) the effective axis alignment (the “relative orientation” of manifolds) (Fig. S11a). We found that the difference in manifold capacity between Naïve and Trained fish can be attributed primarily to a decrease in the effective radius and dimension while contributions of covariance patterns (center and axis alignment) were minor (Fig. 3m, Fig. S11b). Hence, training enhanced the separability of representational manifolds by increasing their compactness and decreasing their dimensionality. These geometrical features also accounted for variations in manifold capacity in other brain areas, including a systematic increase in manifold capacity along successive stages of the ventral visual processing stream (Chou et al., 2025), indicating that effective radius and dimension are relevant geometrical features of representational manifolds in different systems.

We next examined how manifold capacity was affected by repeated odor exposure. Separate point cloud manifolds were constructed from odor responses of each trial and manifold capacity was then quantified for each repetition (Fig. S11a,d-e). In Naïve fish, manifold capacity appeared higher initially but no statistically significant trend was detected over repetitions. Likewise, no significant trends were found for effective radius or dimension. In Trained fish, in contrast, manifold capacity increased consistently and significantly with repeated odor stimulation while the effective radius and dimensionality decreased (Fig. S11d). These trends were qualitatively similar for familiar and unfamiliar odors (Fig. S11e). By contrast, the capacity of point cloud manifolds representing the same repetitions across odors did not differ significantly between training groups (Fig. S11a,f). Further time-resolved analyses showed that manifold geometry increased and decreased abruptly at the onset and offset of odor presentation, respectively, but remained relatively stable during the long tonic phase of the odor response (Fig. S11g). These observations indicate that repeated odor stimulation resulted in more efficient untangling of odor representations in Trained fish.

### 2.5 Temporal structure of odor responses

To further examine the temporal structure of odor representations we determined correlations between activity evoked by repetitions of the same odor in overlapping 300 ms time bins (Fig. 3n). Prior to odor onset, no difference in trial-to-trial correlations was detected between Naïve and Trained fish. Shortly after odor onset, correlations between trials of the same odor increased and approached a plateau after a first peak. After odor offset, correlations peaked again and returned to baseline levels. In Trained fish, trial-to-trial correlations were significantly higher throughout most of the odor response and following odor offset. These differences were most pronounced during the peak phases and consistent for both familiar and unfamiliar odors (Fig. S9e). Hence, odor representations in Trained fish were more reliable throughout the odor response, particularly following changes in odor concentration.

In Trained fish, trial-to-trial correlations increased significantly faster after stimulus onset and peaked earlier than in Naïve fish (Fig. 3o), both in response to familiar and unfamiliar odors. These observations were quantified by linear and exponential fits (Fig. S9f-h). Hence, odor representations were both more reliable and more dynamic in Trained fish, which may enable more efficient odor tracking.

### 2.6 Decomposition of novelty and identity components

The finding that response attenuation is not correlated to response intensity or selectivity (Fig. 1k, Fig. 2i, Fig. S4) suggests that novelty and identity may be represented by distinct components (modes) of activity across the same population of neurons. To test this hypothesis we decomposed activity patterns using demixed principal component analysis (dPCA), a method to factorize activity patterns across mixed-selectivity neurons into modes representing defined task or environmental (external) variables (Kobak et al., 2016).

Activity patterns were decomposed using odor identity and time within a trial as external variables. This decomposition yields different sets of demixed principal components (dPCs) representing information primarily about one external variable (“odor” or “time”) or their combination (“odor x time”) (Fig. 4a). We surmised that if novelty and identity are represented by different modes, these modes should both be contained among dPCs representing “odor” or “odor x time”, which are together referred to as *stimulus* dPCs. Among *stimulus* dPCs, novelty-related dPCs are expected to show low odor specificity and to decline over repetitions while identity-related dPCs should be more odor-selective and stable. *Time* dPCs, on the other hand, are assumed to contain information that is not closely related to odor identity and novelty.

**Figure 4.**
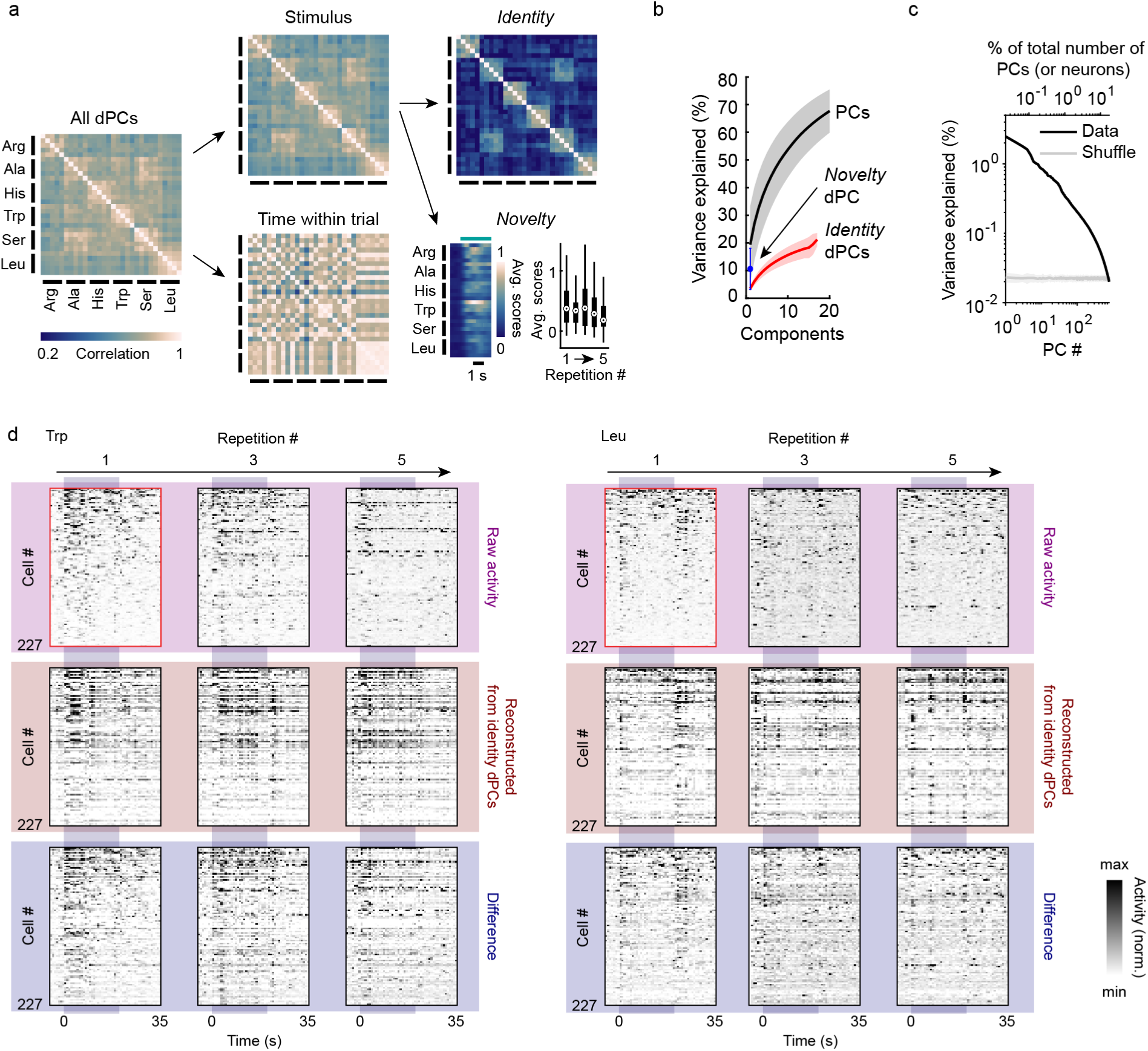
Decomposition of population activity using dPCA. **a** Workflow of dPC decomposition, illustrated by average inter-trial correlations computed from the corresponding dPC scores in Naïve fish. **b** Cumulative percentage of total variance explained by identity dPCs and novelty dPCs, compared to the first 20 classical PCs. **c** Percentage of total variance explained by classical PCs on pooled data from all Naïve fish. The top ∼20% PCs individually explain more variance than loading-shuffled PCs. Linear decompositions therefore explain only a small fraction of the variance, indicating a high embedding dimensionality. **d** Example dPCA decomposition of responses to three repetitions of two single odors (Left: Trp; right: Leu) in the same Naïve fish. In all plots, cells are sorted by decreasing mean firing rate during repetition 1 (outlined in red; color maps normalized individually for each plot). Top: full response (“raw activity”; no dPC decomposition). Note that activity changes substantially after the first repetition. Middle: activity reconstructed from identity dPCs. Note that responses are more consistent across repetitions. Bottom: difference between native and reconstructed activity. This residual activity includes mostly identity-insensitive components, including the novelty component. Note that, while Leu was introduced later in the experiment, observations are qualitatively similar.

In datasets from each fish we selected the 20 dPCs contributing the highest fraction of total variance. Individual dPCs typically represented combinatorial activity across multiple neurons (Fig. S12c). We then computed correlations between activity patterns reconstructed from the time-averaged contributions (scores) of *stimulus* or *time* dPCs for each trial (Fig. 4a). As expected, odor-specific structure was prominent in correlation matrices constructed from *stimulus* but not *time* dPCs (Fig. 4a). We thus retained only *stimulus* dPCs for further analyses, which accounted for 15 – 18 of the 20 selected dPCs in each fish.

We consistently observed that the *stimulus* dPC representing the most variance made strong contributions to all responses, and that these contributions decreased over repetitions. Removing this dPC from the set of *stimulus* dPCs enhanced odor-specific structure in correlation matrices by removing a global positive offset in correlation (Fig. 4a). We therefore defined this dPC as the *novelty* mode and the remaining stimulus dPCs as *identity* modes. On average, these modes accounted for 10.6% and 21.2%, respectively, of the total variance (Fig. 4b). Visualizing the contribution of the *identity* modes to overall activity patterns (Fig. 4d) confirmed that activity reconstructed from *identity* modes was more stable than the overall activity, which changed and decreased substantially over repetitions.

Unlike standard principal components, dPCs are not strictly orthogonal by design. Nonetheless, pairwise correlations between dPCs were narrowly distributed around zero (Fig. S12d-e). Hence, different modes represented largely non-redundant components of the observed population activity. Loadings of dPCs were broadly distributed across odors with no obvious clustering (Fig. S12a-c), demonstrating that modes did not map onto individual odors but represented patterns of activity across mixed-selectivity neurons.

The finding that multiple near-orthogonal *identity* modes contributed similar amounts of variance (Fig. 4b) suggests that odor-evoked activity in pDp is high-dimensional. Consistent with this hypothesis, the number of non-demixed principal components whose variance exceeded chance levels, as determined by shuffling of weights, was usually >10% of the total number of PCs (or >10% of the number of recorded neurons), both in data from individual fish and in pooled data (Fig. 4c).

We next asked how *novelty* and *identity* modes were affected by experience. The *novelty* dPC had larger amplitude in Trained fish and decayed more prominently and consistently over repetitions than in Naïve fish (Fig. 5a-c). To further analyze *identity*-related response components we first compared correlations between the time-averaged contributions of *identity* modes, or between activity patterns reconstructed from these modes. Differences between the first and subsequent repetitions of each odor (Fig. 5d) were substantially reduced compared to differences between the original correlations (full activity vectors; Fig. 3e), confirming that *identity* modes contained little novelty-related activity. Removing activity reconstructed from *identity* modes from the original activity patterns by subtraction abolished most odor-specific structure in correlation matrices (Fig. 5d). Furthermore, trial-to-trial correlations (same odors) between activity patterns reconstructed from *identity* modes were substantially higher than trial-to-trial correlations between original activity patterns, consistent with a selective enrichment of odor-specific activity (Fig. 5e, Fig. S15f). While activity reconstructed from *identity* modes still decayed over trials (Fig. 5f,g), this decay was less pronounced than in *novelty* modes (Fig. 5b,c). These results further confirm that novelty and identity are represented by largely independent modes of population activity, both in Naïve and Trained fish.

**Figure 5.**
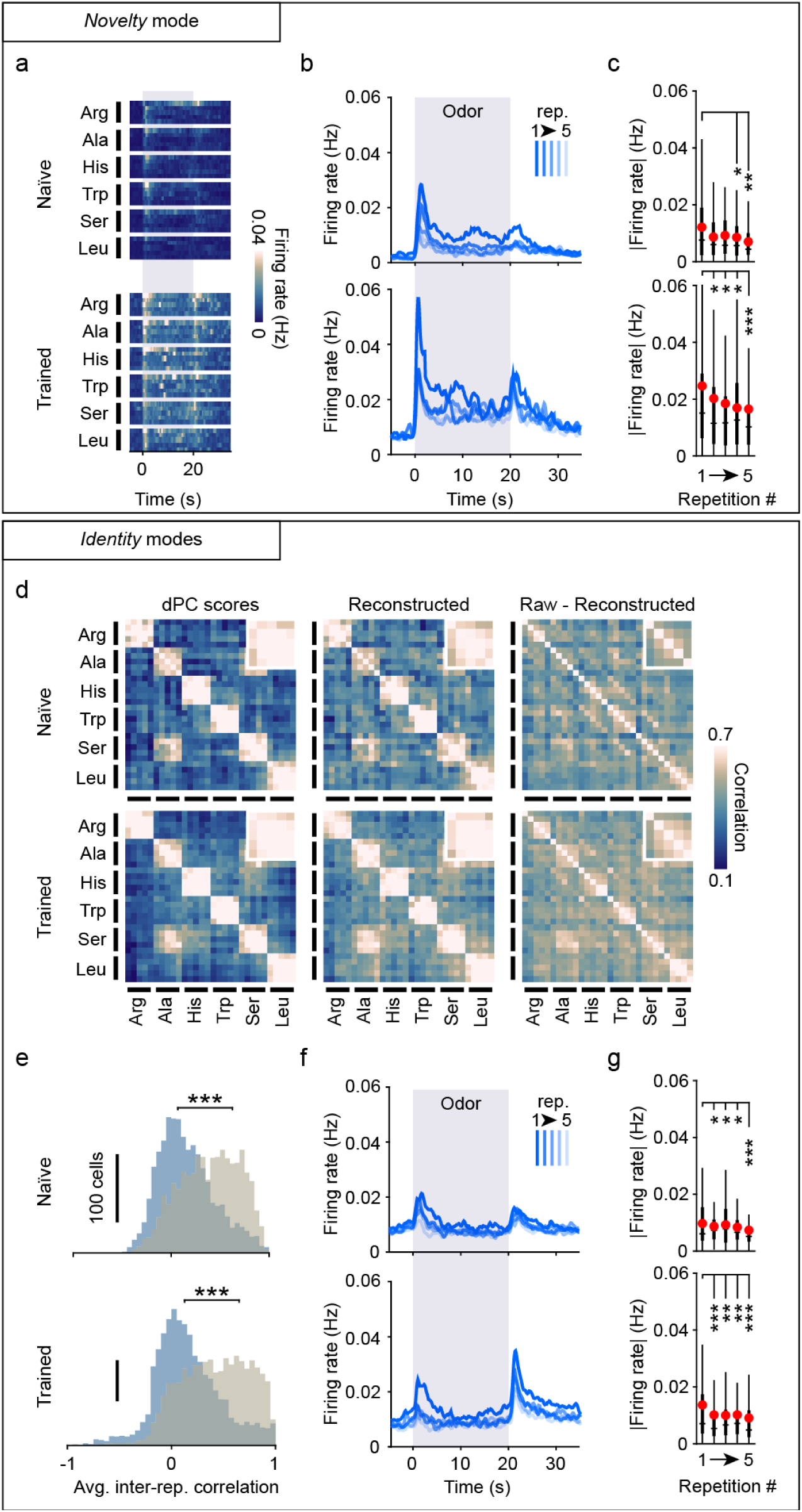
Reconstruction of activity from identity and novelty-associated modes. **a** Time course of average firing rates reconstructed from the novelty dPC only, across repetitions of each odor. Top: Naïve fish; bottom: Trained fish. **b** Average time course of reconstructed activity from novelty dPC in successive repetitions. **c** Average firing rates reconstructed from novelty dPC as a function of repetition number in Naïve (top) and Trained (bottom) fish. Paired Wilcoxon signed rank tests between |Firing rate| on repetition 1 and other repetitions. We use |Firing rate| instead of firing rate because reconstructed activity is not strictly positive. **d** Inter-trial correlations based on identity dPCs, across groups. Left: correlations based on dPC scores. Middle: correlations based on reconstructed cellular activity. Right: correlations based on difference between measured and reconstructed activity (raw – reconstructed). Insets: correlations across same-odor repetitions. **e** Average single-cell tuning correlation across repetitions of the same stimulus, based on raw activity (purple) vs reconstructed activity from identity dPCs (beige). Top: Naïve fish; bottom: Trained fish. Mann-Whitney U test, p(Naïve) = 6.98·10^-161^, p(Trained) = 1.42·10^-280^. **f-g** Same as **b-c** but using activity reconstructed from the identity dPCs. Box plots: 25^th^ and 75^th^ percentile; whiskers: lowest and highest non-outlier values; horizontal line: median; red circle: mean. *: p*<*0.05; **: p*<*0.01; ***: p*<*0.001.

### 2.7 Representation of temporal context

We next examined whether the evolution of odor representations across trials exhibits an underlying global organization related to the progression of time. We first quantified the direction and magnitude of representational changes between each repetition by vectors representing the difference between successive time-averaged odor response patterns (“difference vectors”) and measured their correlation between odors (Fig. 6a, top). Vector correlations were small but significantly greater than zero (Fig. 6a), indicating that representational changes may include a consistent component.

**Figure 6.**
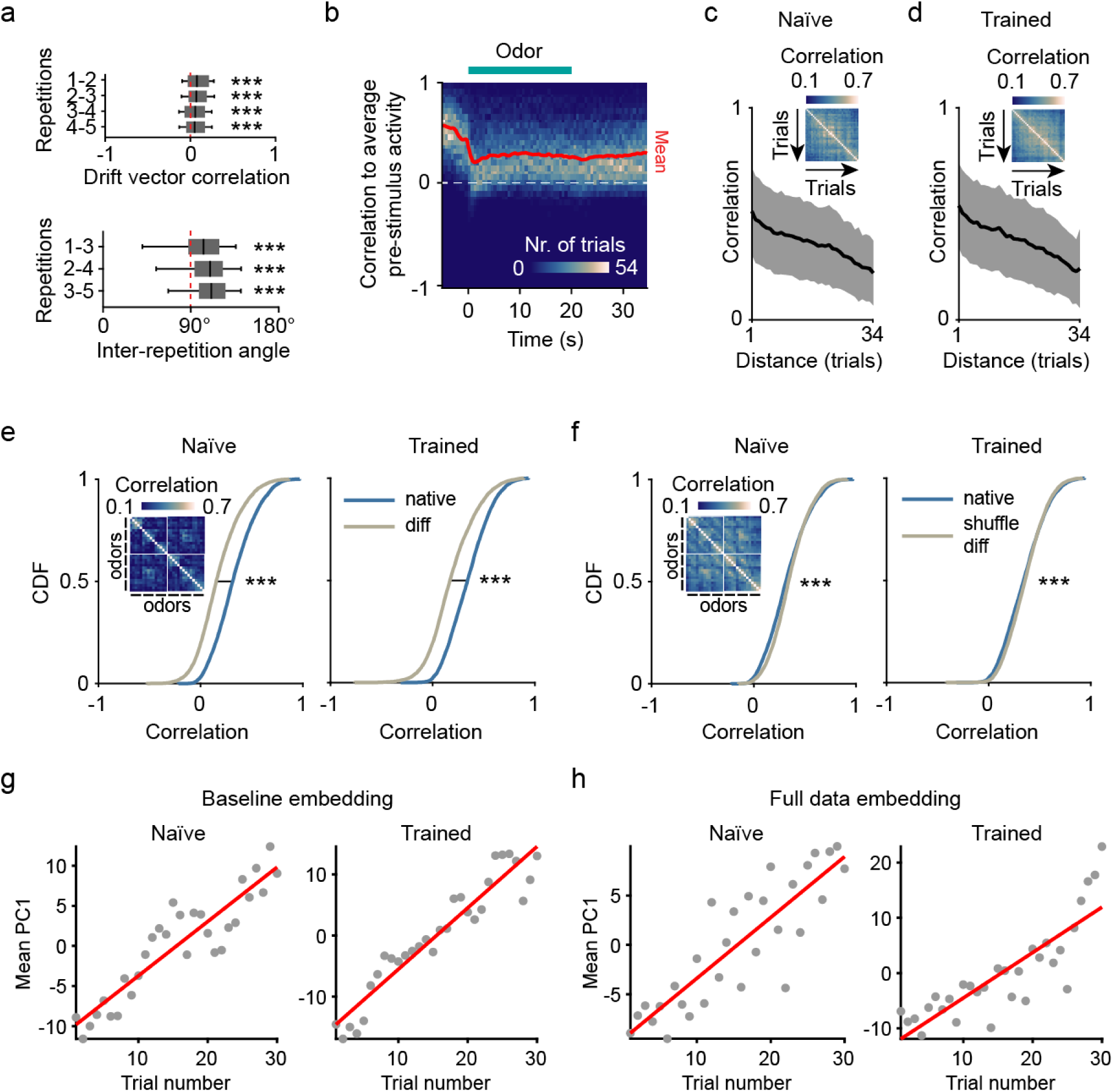
Baseline drift. **a** Top: Correlation of N-dimensional drift vectors (connecting the mean response patterns to consecutive same-odor repetitions) in all fish. For each repetition pair, the median vector correlation was significantly larger than zero (n = 390, 405, 452 or 479 vectors, respectively. In all cases, p < 10^-5^ with 10^6^ bootstrap resamples). Bottom: Triplet angles across repetitions. The median was consistently >90° (in each case: n = 210 angles, p < 10^-6^ with 10^6^ bootstrap resamples), implying that inter-repetition drift not independent. Box plots: 25^th^ and 75^th^ percentile; whiskers: mean ± SD; vertical line: median. **b** Sliding-window histogram of correlations between activity in successive time windows (duration: 300 ms) and pre-stimulus activity (mean over 20 s), averaged across trials and fish (Naïve only). Correlations decrease upon odor stimulation but remain positive, indicating that pre-stimulus and response activity are not orthogonal (see also Fig. S10). Red line: bin mean. **c-d** Inter-trial correlation of pre-stimulus (baseline) activity patterns, with respect to temporal distance, in Naïve (n = 14 fish, **c**) and Trained (n = 21 fish, **d**) fish (mean ± SD). Insets: inter-trial correlation between time-averaged pre-stimulus activity vectors. Trials are sorted chronologically. **e** Cumulative distribution of all pairwise inter-trial correlations between raw and baseline-subtracted response vectors in Naïve (left) and Trained (right) fish. Mann-Whitney U test: median(Naïve) = 0.153, p(Naïve) = 4.00·10^-293^, median(Trained) = 0.169, p(Trained) < 2.2·10^-308^. See also Fig. S13a. **f** Same as **e** but after subtraction of cell-shuffled baseline patterns, which slightly increased correlations. Mann-Whitney U test: median(Naïve) = 0.331, p(Naïve) = 3.0·10^-3^, median(Trained) = 0.351, p(Trained) = 9.79·10^-4^. See also Fig. S13b. **g** Mean PC1 scores of pooled pre-stimulus activity from Naïve (left) and Trained (right) fish across trials (see also Fig. S15a). Red line: linear regression. Pearson correlation values: r(Naïve) = 0.912, p(Naïve) = 2.46·10^-12^; r(Trained) = 0.951, p(Trained) = 8.37·10^-16^. **h** Same as **g** but using full response data embeddings. Pearson correlations: r(Naïve) = 0.861, p(Naïve) = 1.01·10^-9^; r(Trained) = 0.843, p(Trained) = 4.85·10^-9^. ***: p<0.001.

We next examined whether responses to the same odors changed in consistent directions between successive repetitions. If directional changes between repetitions are random, successive difference vectors should be nearly orthogonal given the high-dimensional embedding space. However, angles between successive difference vectors were consistently >90° (Fig. 6a, bottom), indicating that changes were partially aligned across repetitions. These observations suggest that the evolution of odor responses across repetitions involves a common underlying drift process.

Given that pDp neurons showed significant activity prior to odor stimulation in vivo, a common contribution to odor representations may originate from odor-independent patterns of baseline activity. To test this hypothesis, we first examined the correlation of population activity in a sliding 300 ms time window to the time-averaged pre-stimulus (baseline) pattern. Correlations to the baseline pattern were high (∼0.5) prior to odor stimulation and decreased at odor onset. However, correlations remained substantially and significantly above than zero throughout the odor response and beyond (Fig. 6b). Hence, activity during baseline persists, in part, during the odor response.

We next averaged activity over 20 s prior to each stimulus to capture patterns of baseline activity and analyzed their correlations across trials. In Naïve fish, baseline correlations were high between successive trials (r = 0.53, n = 406 trial pairs across 14 Naïve fish) and decreased nearly linearly with increasing temporal distance at an average rate of 0.17/h, thus dropping from 0.53±0.05 to 0.22±0.04 (mean ± s.e.m.) over the course of the experiment (Fig. 6c). Similar rates of change were observed in Trained fish, with correlations falling from 0.54±0.04 to 0.24±0.04 (mean ± s.e.m., equivalent to an average of 0.18/h Fig. 6d). Hence, baseline activity drifted over time on a timescale of minutes to hours. Further analyses confirmed that this drift is a population phenomenon that cannot be attributed to a small number of outlier neurons, and that it cannot be explained by artifacts such as mechanical instability of the tissue (Fig. S14). Hence, baseline activity exhibited a constant drift that may contribute to the coherent change in odor representations over repetitions.

To examine the contribution of baseline activity to odor responses we assumed that baseline and odor-evoked activity are independent and interact additively. To test this hypothesis we subtracted time-averaged baseline activity (20 s) from the subsequent odor response in each trial. If interactions are additive, baseline subtraction should globally decrease pattern correlations by removing a common pattern component. Alternatively, baseline subtraction should not decrease or even increase pattern correlations because baseline subtraction imports correlated activity into odor responses. We found that pattern correlations were substantially lower after baseline subtraction, implying that subtraction removed a common component from population activity patterns (Fig. 6e). By contrast, subtracting a cell-shuffled baseline slightly increased correlations, consistent with an import of correlated activity (Fig. 6f, Fig. S13b). Hence, activity during odor presentation can be described, in first approximation, by the additive combination of an odor-independent baseline and an odor-specific response pattern (Fig. 6e, Fig. S13a). Baseline activity therefore makes a substantial and specific contribution to the overall activity during odor presentation.

The first response remained noticeably different from subsequent responses even after subtraction of baseline activity, as revealed by same-odor correlations across repetitions both in Naïve and in Trained fish (Fig. S13a, inset). The contribution of drifting baseline activity can therefore not account for the initial novelty-related changes in odor representations.

We next examined whether baseline drift exhibits a directional bias. If so, the drift process should be captured, in first approximation, by a linear projection axis. Indeed, projecting baseline activity onto the first (non-demixed) principal component (PC1) revealed a high correlation between PC1 and trial number (Fig. 6g). This correlation was only slightly lower when PC1 was computed from activity during odor presentation (Fig. 6h). These observations further show that population activity in pDp contains a prominent component that encodes temporal context independent of the presence or absence of an odor.

Consistent with the additive interaction between baseline and odor-evoked activity, subtraction of the baseline component strongly reduced the correlation between PC1 and trial number (Fig. S13c). Similar observations were made across Naïve and Trained fish, both in pooled data and in individual fish (Fig. S13d). These results further support the conclusion that a slowly drifting component of population activity in pDp contains information about temporal context.

### 2.8 Low-dimensional embeddings of population activity

Our results indicate that population activity in pDp is high-dimensional and represents multiple categories of information including stimulus novelty, stimulus identity and temporal context by distributed activity across mixed-selectivity neurons. To further understand the organization of this activity we performed low-dimensional embeddings using linear (PCA) and non-linear (UMAP) approaches. Sequences of activity patterns during odor presentation in each trial are thus visualized by trajectories in the low-dimensional embedding space.

In the space defined by the first two principal components, trajectories measured in Naïve fish overlapped extensively (Fig. 7a; colors represent odors; line thickness represents repetition number). Nonetheless, trajectories representing different odors were biased towards different territories. Moreover, trajectories representing later repetitions (thinner lines) were systematically associated with higher values along the PC1 axis, consistent with the close relationship between trial number and PC1 extracted from time-averaged activity patterns (Fig. 6h, Fig. S13d). In Trained fish, individual trials and groups of trials representing different odors were more clearly separated from each other. This was evident particularly along PC2, while PC1 represented trial number (time). In both Naïve and Trained fish, trajectories representing Leu (yellow) were shifted to higher values along the PC1 axis, reflecting the fact that Leu was introduced late in the experiment.

**Figure 7.**
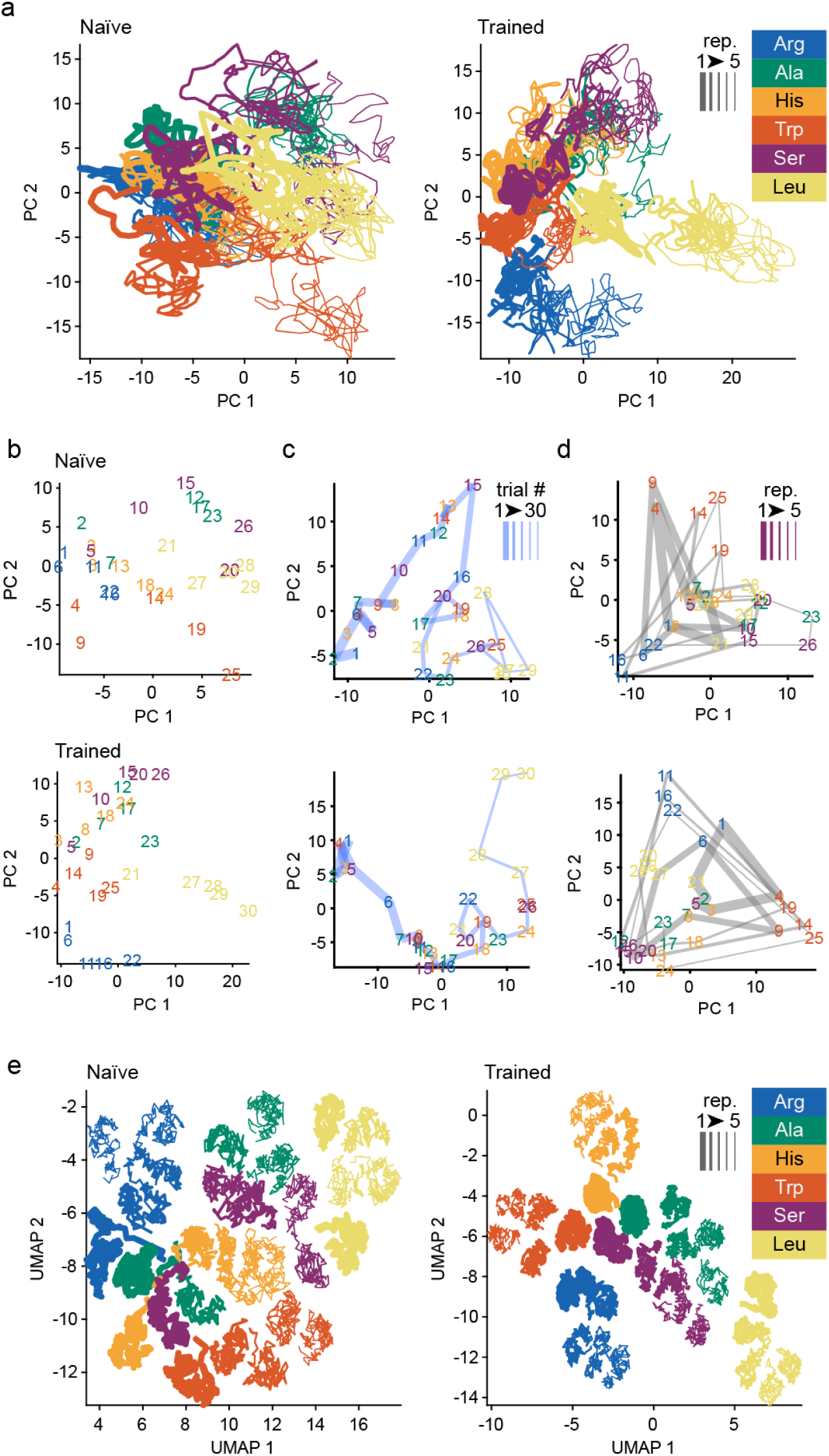
Low-dimensional embedding of population dynamics. **a** First 2 PCs of pooled responses from Naïve and Trained fish. Colors indicate different stimuli, lines connect patterns in successive time bins, line thickness indicates repetition number. **b** Same as **a** but individual trial centroids are displayed by their respective trial number. Colors indicate stimulus identity. Top: Naïve. Bottom: Trained. **c** Trial number embedding (as in **b**) of pooled pre-stimulus activity. The blue curve connects consecutive trials and becomes thinner over trials. PC axes are computed from baseline data and therefore different from **a-b** (see Fig. S16 for the same data, projected onto PC axes from **a-b**). **d** Trial number embedding (as in **b**) of pooled baseline-subtracted response data. Gray lines connect the same repetition number across stimuli; thickness corresponds to repetition number. The order of connection among stimuli is chosen to maximize polygon convexity within each repetition. Note that PC axes are computed on baseline-subtracted data and therefore different from **a-c**. **e** UMAP embedding of pooled responses.

To further simplify the visualization of activity patterns we represented the centroids of trajectories by their corresponding trial numbers, colored by odors (Fig. 7b). These visualizations confirmed the representation of time by PC1 (increasing trial numbers along the PC1 axis) and more consistent clustering of trajectories by odor (colors) in Trained fish. We next projected centroids of trajectories representing 20 s of baseline activity prior to the presentation of each odor into the same embedding space (Fig. S16) or onto the first two PCs computed from baseline activity (Fig. 7c). Again, trial number increased systematically along PC1, even when PC1 was derived from activity during odor presentation. These results confirm the presence of a prominent component of population activity that is largely independent of odor presentation and represents temporal context.

Finally, we subtracted baseline activity from odor-evoked activity in each trial and embedded the resulting activity patterns in the space of the first two PCs (newly computed after baseline subtraction) (Fig. 7d). As expected, baseline subtraction abolished the systematic relationship between trial number and PC1. We next connected centroids representing trials of the same block by lines to visualize the territory of the embedding space that was spanned by representations of different odors in each block. With increasing block number, the size of the territory increased, indicating that the variance represented by the first two PCs was used more extensively for the representation of different odors. This effect was prominent particularly in Trained fish.

In UMAP embeddings (Fig. 7e), trajectories representing different trials and odors were separated better than in PCA embeddings. The separation of trajectories and their clustering by odors was again more prominent in Trained fish. Time (trial number) was not aligned with a global UMAP axis but trajectories within each odor cluster were systematically organized by repetition number. Further low-dimensional embeddings confirmed that baseline activity represented the progression of time in the absence or presence of odors (Fig. S15). In summary, low-dimensional embeddings therefore confirmed and extended conclusions derived from other analyses.

### 2.9 Phenomenological model

Our results indicate that odor-evoked activity in pDp can be decomposed into at least three components: (1) a novelty component that decreases with repeated stimulation, (2) an odor identity-dependent component that becomes more prominent with repeated odor stimulation, and (3) a time-dependent component that can be attributed largely to slowly drifting baseline activity. To further explore to what extent these components can account for experimental observations we developed a simple, purely phenomenological model. Odor responses were modeled by three additive terms representing an odor-specific identity component, a novelty component, and an odor-independent drift component. While the novelty term decreased rapidly over repetitions, the identity component increased slowly and the drift component rotated and decayed even more slowly over trials. Each component was represented by a prescribed direction in neural state space, with biologically realistic lognormal firing rate distributions and spatio-temporally correlated noise. This model was contrasted with a null model representing stimulus identity along static and orthogonal population modes (Fig. 8a-c).

**Figure 8.**
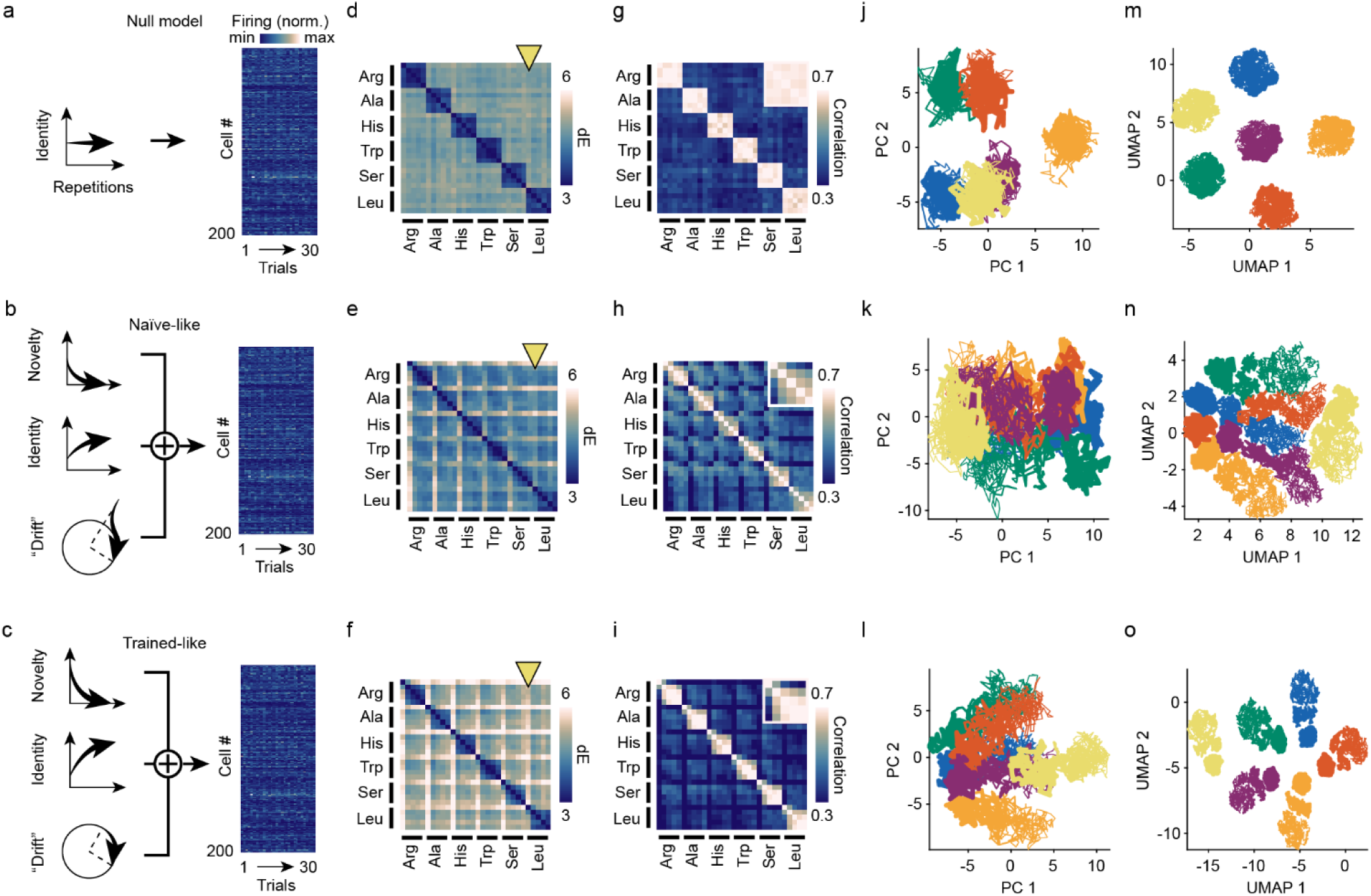
Phenomenological model. **a-c** Schematic representation of the main components of the ‘null model’ (**a**), ‘Naïve-like’ (**b**) and ‘Trained-like’ (**c**) implementations. In the null model, odors were represented by orthogonal and invariant activity patterns, similar to the expected output of a classical autoassociative classifier. In the Naïve-like and Trained-like models, three time-varying linear components add up to one activity pattern per trial. The ‘novelty’ component decays independently over repetitions of each odor while the ‘identity’ component increases over repetitions. The “drift” component represents baseline activity, contributing to the odor response pattern and drifting over trials at a constant rate. Population firing patterns are simulated by adding biologically realistic neuronal offsets and noise. Motivated by our observation that the attenuation of odor responses over trials is less odor-specific in Naïve fish, the initial magnitude of the novelty component and the asymptotic magnitude of the identity component were lower in the Naïve-like model (**b**). **d-o** Qualitative results from the three models. Datasets were initialized using the same 5 random seeds across model types; therefore, differences are only due to the choice in geometric parameters across models. **d-f** Average inter-trial Euclidean distances. **g-i** Inter-trial correlations. Inset: correlations across same-odor repetitions. **j-l** First 2 PCs of pooled data. Colors indicate different stimuli and line thickness indicates repetition number (higher repetition number: thinner line). **m-o** 2D UMAP of pooled data. **d,g,j,m** Results from the Null model. **e,h,k,n** Results from the Naïve-like model. **f,i,l,o** Results from the Trained-like model.

Effects of training were simulated by modest changes in model parameters reflecting experimental observations: in the model representing Trained fish (“Trained-like”), amplitudes of the identity and novelty components were increased while the amplitude and decay of the baseline contribution were decreased relative to the “Naïve-like” model. Without systematic parameter variations, the models reproduced basic observations including the decrease in Euclidean distances between odor representations after the first repetition (Fig. 8e-f), the representation of novelty information (repetition number) along the first principal component, and the segregation of odor identity along the second principal component (Fig. 8k-l). By contrast, none of these observations were reproduced by the null model (Fig. 8d,g,j,m).

Consistent with experimental observations, response amplitudes and, consequently, Euclidean distances were higher in the Trained-like than in the Naïve-like model. Moreover, novelty was represented more distinctly, as seen by the higher Euclidean distance between the first and subsequent repetitions, particularly for the odor that was introduced late in the experiment (Leu). Low-dimensional embeddings showed a more consistent separation of activity trajectories in the Trained-like model. Hence, the odor-specificity of the novelty response was enhanced (Fig. 8e-f) and representations of different odors were more distinct in the Trained-like model (Fig. 8h-i,k-l,n-o). Effects of training could therefore be reproduced by parameter changes that result in more distinct representations of odor identity and novelty relative to baseline activity. These observations confirm the conclusion that pDp jointly represents multiple olfactory and temporal stimulus features by largely independent components of population activity, and that learning enhances the accessibility of odor-related information by modifying neural manifolds. However, as the model is purely descriptive, it cannot provide insights into the underlying mechanisms.

## 3 Discussion

Lateral optical access through a prism allowed us to measure odor-evoked population activity in pDp of adult zebrafish in vivo. Population activity in pDp contained information not only about odor identity but also about novelty and temporal context. Discrimination training increased the reliability of odor responses, accelerated response dynamics, and enhanced the separability of representational manifolds. These results reinforce the notion that pDp continuously updates an internal representation of odor space according to an individual’s experience, rather than storing information about distinct odors in an itemized memory (Hu et al., 2026).

The experience-driven untangling of representational manifolds, as well as the changes in pattern correlation and response dynamics, are likely to support rapid and generalizable odor classification by facilitating access to relevant information. Moreover, we found that odor identity is represented together with information about stimulus history (novelty) and temporal context by mixed-selectivity neurons. Hence, pDp integrates representations of multiple stimulus features including time, which is critical for episodic memory and supports generalization (Rigotti et al., 2013; Posani et al., 2026). Together, these findings reframe pDp as a sensory context map that is updated in an experience-dependent manner by geometrical modifications of representational manifolds.

### 3.1 Odor-specific novelty detection in pDp

Odor responses in pDp were distributed, stimulus-specific yet variable, and followed a phasic-tonic time course, consistent with previous observations in an ex-vivo preparation of the zebrafish brain (Yaksi et al., 2009; Blumhagen et al., 2011). In-vivo measurements further confirmed that odor representations are attenuated and modified by repeated stimulation, particularly after the first presentation of an odor (Jacobson et al., 2018). Using interspersed applications of multiple similar odors we found that this novelty-related plasticity exhibits considerable odor-selectivity. Neuronal circuits in pDp therefore function as odor-selective novelty filters on timescales of (tens of) minutes.

As novelty filters emphasize unpredicted inputs they may support the detection of salient and potentially meaningful events. Computationally, novelty detection is closely related to predictive processing and can provide an error signal to update internal models (Rao and Ballard, 1999; Clark, 2013). In classical models, novelty filtering can be implemented by structured negative feedback reflecting previous inputs, which in turn can be established by activity-dependent plasticity of recurrent connectivity (Oja, 1982; Kohonen, 1984; Chklovskii et al., 2004; Vogels et al., 2011; Schulz et al., 2021). In principle, the underlying mechanisms of novelty detection are consistent with the general architecture of autoassociative memory networks and with the observed odor-selective attenuation of responses in pDp (Kohonen, 1984).

### 3.2 Organization of multimodal representations in pDp

Multiple observations—including the rapid pattern changes at stimulus transitions and the generalization of experience-dependent effects across similar odors—confirm the conclusion that odor-evoked activity in PCx/pDp does not show obvious signs of convergent attractor dynamics (Franks et al., 2011; Roland et al., 2017; Pashkovski et al., 2020; Schoonover et al., 2021; Hu et al., 2026). Our results thus reinforce the view that PCx/pDp maintain flexible representations of odor space that are defined by the geometry of representational manifolds. pDp operates in a state of precise excitation-inhibition balance (Rupprecht and Friedrich, 2018), which may account, in part, for the observed variability of neuronal activity. In such networks, structured connectivity can confine activity onto manifolds that encode information reliably and efficiently even though the activity of individual neurons appears unstructured (Denève and Machens, 2016; Denève et al., 2017; Meissner-Bernard et al., 2025b).

As activity of pDp neurons contained information not only about odor identity but also about novelty and temporal relationships, pDp contains joint multimodal representations of olfactory and non-olfactory stimulus features related to time. These stimulus features could be assigned to different modes of population activity, indicating that they are, in first approximation, represented along different state space dimensions.

In vivo activity measurements revealed a prominent odor-independent, drifting component of population activity that contained information about the progression of time over minutes to hours. This timescale is distinct from representational drift in PCx and hippocampus over days and weeks (Rubin et al., 2015; Schoonover et al., 2021; Keinath et al., 2022), indicating that the underlying processes are likely different. It remains to be determined whether drift in pDp is independent of odor stimulation, reflecting elapsed time, or whether it is modulated by sensory input, reflecting episodic time. A drift in population activity on a timescale of seconds to hours was observed in the lateral entorhinal cortex of rodents (Tsao et al., 2018; Kanter et al., 2025). As observed in pDp, this drift is superimposed onto activity representing other information, raising the possibility that representations of time in these brain areas are functionally related.

### 3.3 Modification of multimodal olfactory representations by long-term experience

After training of fish in an odor discrimination task, activity in pDp did not represent specific associations between odors and rewards. These observations further support the conclusion that pDp does not assign odors to discrete, task-dependent classes by pattern completion. Moreover, they indicate that pDp contains no explicit representation of valence. An experience-dependent joint representation of odor space and valence has been observed in an adjacent telencephalic area (Frank et al., 2019), supporting the hypothesis that the separation of neural manifolds in pDp preceeds the flexible association between odors and valence in higher brain areas.

Consistent with previous observations (Hu et al., 2026), discrimination training enhanced odor responses and their separability. While effects of training on odor classification were relatively weak when assessed by standard classifiers, pronounced and consistent increases in the separability of neural manifolds were observed using GLUE theory (Chou et al., 2025). This observation further reinforces the conclusion that pDp mediates representational learning rather than direct pattern classification. GLUE theory directly quantifies the “untangledness” of potentially complex point cloud manifolds without averaging, and it links classification to manifold geometry. GLUE analysis revealed that experience significantly enhanced the separability of manifolds primarily by increasing their “compactness” and by decreasing their effective dimensionality. These geometrical features have recently been found to be main determinants of manifold capacity also in other systems (Chou et al., 2025; Hu et al., 2026). Additional time-resolved analyses of trial-to-trial correlations showed that odor responses were more reliable in Trained fish, particularly after odor onset and offset. Together, these results reveal experience-driven changes in neural manifolds that support the classification of familiar and related odors.

Further results show that experience increased the magnitude and stimulus-specificity of the novelty-related response component. As a consequence, repeated odor stimulation enhanced the odor-selectivity of population responses more prominently in Trained than in Naïve fish. These observations further support the notion that experience-dependent modifications of representational neural manifolds support perceptual learning and distributed learning processes across brain areas (Haberly, 2001),. In summary, our results show that representational manifolds in pDp integrate information about odor identity, novelty and time. These representations are modified by experience to enhance the accessibility and reliability of potentially relevant information. As novelty and temporal relations are critical for episodic memories, pDp may thus contribute to the formation of episodic memory by establishing joint representations of olfactory and temporal information.

## 4 Methods

### 4.1 Experimental subjects

All experiments were performed on adult, 6-12 months old zebrafish (*Danio rerio*) without selecting for sex. Data from both sexes was used in approximately equal proportion and combined, as the variable of sex is not considered in this study. Fish were raised and kept under standard laboratory conditions (24-28°C; 13 h light cycle). All experimental protocols were approved by the Veterinary Department of the Kanton Basel-Stadt (Switzerland).

Lesioning experiments were performed on tg(vGlut1:eGFP)^+/-^ fish, which express enhanced green fluorescent protein in a large and distributed subpopulation of glutamatergic neurons. This line was used because the pervasive green background fluorescence in the brain simplified post-mortem assessment of mechanical damage to the forebrain with confocal microscopy.

Calcium imaging experiments were performed in tg(NeuroD:GCaMP6f) fish (Rupprecht et al., 2016), which express the genetically encoded calcium indicator GCaMP6f (Chen et al., 2013) in almost all forebrain neurons, except in the olfactory bulb (Fig. S1c). Fish were bred to have the leucistic *nacre* pigmentation (Lister et al., 1999) to reduce light absorption.

Anaesthesia was performed as follows: first, the fish were individually placed into 0.24 g/L MS222 (induction bath) until unresponsive to a light tail pinch, then quickly intubated with 0.074 g/L MS222 under constant perfusion through a cannula in the mouth. In a subset of experiments, fish were immersed in 5 mg/L Lidocaine prior to anaesthesia. Drugs were dissolved in housing system water (except for post-operative analgesia prior to 2-photon imaging, which was dissolved in ACSF) at pH 6.5-7, reflecting housing conditions.

### 4.2 Behavioral training for the lesioning experiment

Behavioral training was carried out as described (Namekawa et al., 2018). Briefly, adult zebrafish were first habituated individually to the training environment for ∼36 h without food. Subsequently, training was carried out for 5 consecutive days under constant water exchange. Throughout habituation and training, fish were kept in water from the same system they were originally housed in. Each animal received 18 odor applications (trials) per day (9 CS+ and 9 CS-presentations). Odors were introduced into the perfusion water for 30 s with minimal disturbance of the flow rate. Application of the CS+ but not the CS-was followed by delivery of a small amount of fish food (crushed flakes) was delivered into a feeding ring at a specific location. The inter-trial interval was 20 minutes. The behavioral performance of the fish was analyzed daily, such that “putative learners” could be identified by the end of training day 5, using data from days 1-4. At the end of training day 5, putative learners underwent either the lesioning or a sham surgical procedure, after which they were returned to the training setup to recover overnight. On day 6, discrimination performance was tested following the standard behavioral protocol. Subsequently, fish were euthanized within 24 h, and the whole brain was extracted and post-fixed for 24 h at 4°C for histological analysis.

### 4.3 Lesioning of the pallium

To avoid experimenter bias based on behavioral observations, the individual identifier for each pre-trained fish was first substituted with a unique sham identifier by a different person, who also held the mapping key. This key was only revealed after the surgical procedure and used to correctly return each animal to its holding tank in the behavioral setup. To lesion the pallium, we modified a previously established transcranial infusion protocol (Zhou et al., 2014; Satou et al., 2022). Each animal was first anaesthetized as described above, quickly immobilized into a custom sleeve holder and intubated. After gentle removal of a small portion of skin above Dp, we opened a small craniotomy to expose the dorsal surface of the pallium. To ablate the region of interest, we gently lowered a glass cannula (i.d. ∼150 µm, o.d. ∼250 µm) until it made contact with the meninges, and applied a single brief pulse of negative pressure. Some localized and temporary bleeding occurred occasionally. Approximately half of the experimental animals received a craniotomy, but no ablation (sham group). To avoid bias on the part of the experimenter, the assignment of fish to the ablated or sham group was made only after opening the craniotomy using a random number generator. The procedure for bilateral ablation of Dp lasted approximately 5 minutes, after which the fish were allowed to recover under observation for 10 minutes. The animals were then returned to the behavioral tanks.

### 4.4 Behavioral training for functional microscopy

Animals for calcium imaging experiments were trained as described above with minor modifications. First, the set of conditioned stimuli was composed of three monomolecular amino acids (L-Arginine, L-Alanine ad L-Histidine, each 6·10^−5^ M), two rewarded (CS+1, CS+2) and one unrewarded (CS-). Reward attribution varied across training groups (Fig. 2e). Second, animals were pre-trained on the same stimulus-reward schedule over 2 weeks in their home tanks in groups of 8 or 9 fish, as described (Hu et al., 2026). The fish were then individually tested for 3 - 5 days in the single-housed setup (Fig. 2f, Fig. S5a-b). Trained groups comprised a total of n = 21 fish.

### 4.5 Calcium imaging

#### 4.5.1 Head fixation and prism placement

Animals were head-fixed as described in (Huang et al., 2020) with minor modifications and placed under the 2-photon microscope in a custom holding tank. A dorsolateral craniotomy was performed over the posterior lateral forebrain (Fig. S1a) to introduce a reflective microprism lateral to the pallium. The prism was glued to a rod and positioned using a motorized manipulator without damaging the brain or the meninges (Fig. S1a). The prism reflected both 2-photon excitation (infrared) and fluorophore emission (visible) light at a 90^◦^ angle (see below), allowing for sagittal plane imaging of the lateral dorsal forebrain up to a depth of 250-300 µm from the lateral surface (Fig. 1a-b, Fig. S1b), covering a large portion of ipsilateral pDp, the nucleus teniae and adjacent areas (Fig. S1b).

After surgery, the animal was placed on the microscope stage and kept in darkness for the duration of the experiment. The liquid inlet targeting the left nostril delivered a constant flow of oxygenated teleost artificial cerebral-spinal fluid (ACSF; 10 ml/min). This was expected to dilute dissolved compounds to half of the initial concentration in approximately 8.3 minutes, according to a first order decay model *dC*/*dt* = −*FR*/*V* · *C*, where C is the concentration, FR is the flow rate, V is the tank’s volume and t is time.

Because an open craniotomy may put the brain in contact with the external medium, fish were continuously held in isotonic ACSF, starting right before the first transcranial incision and until the end of the experiment. Zebrafish are known to be resistant to significant changes in water salt content. The osmolarity of teleost ACSF (∼300 mOsm) is within the range commonly used in fish facilities in general and similar to that of normal housing water. Craniotomized head-fixed fish were only held in ACSF for the duration of the experiment (max. 3 h), and likewise showed no difference in behavior or survival rate, compared to similarly restrained intact animals held in housing system water.

#### 4.5.2 Behavioral tracking during calcium imaging experiment

The fish was illuminated with two 980 nm laser diodes (LDM 0980-050-50) powered using a custom driver and imaged with a FLIR NIR camera (GS2-U3-23S6M-C) and a zoom lens (MVL7000 - 18 108 mm EFL, Navitar) equipped with a 975 nm lowpass filter (Edmund Optics, 86-073). The filter allowed in light from the LEDs, while blocking most light from shorter wavelength sources, including the LSM. The remaining low-intensity background from the 2-photon laser was used to align trial boundaries with the calcium data during post-hoc analysis. Movies were rotated and cropped into three ROIs covering a background area, the head, and the tail. Automated basic analyses of pixel fluctuations in the head and tail ROIs were used to monitor motion of the lip, jaw, operculae or tail. This monitoring ensured that fish did not undergo major changes in behavioral state during the experiment. Moreover, tail motion was quantified (Huang et al., 2020) to assess motor-related neuronal activity (Fig. S3).

#### 4.5.3 Optical setup

GCaMP6f fluorescence was imaged using a custom-build 2-photon microscope based based on a MOM body (Sutter Instruments). GCaMP6f was excited at 930 nm with a Ti:Sapphire laser (Spectra-physics Mai Tai eHP Deep See, 1.8 MW peak power, 80 MHz repetition rate, 70 fs pulse width). The beam was expanded 3 times (AC254-050-B-ML AC254-150-B-ML, Thorlabs) before entering the MOM. XY galvanometer scanners (6215H, Cambridge Technology) scanned the beam across the area of interest. Scan- and tube lens further expanded the beam ∼4 times before entering the objective (XLSLPLN25XSVMP2, Olympus, 25X 0.95 NA, 8 mm WD). Power under the objective was approximately 170 mW. The light cone exiting the objective lens was reflected 90° by the Ag-coated hypothenuse of the microprism (RPB3-1.4-450-1500-AG, OptoSigma, Fig. 1a). Fluorescence emission was again reflected through the prism, then collected by the objective, filtered (ET535/50m, Chroma) and finally projected onto the photomultiplier tube (PMT, H11706P-40 or H7422p-40MOD, Hamamatsu). Image acquisition was controlled using ScanImage 6 (Pologruto et al., 2003, Vidrio Technologies) and MATLAB 2019b (MathWorks).

#### 4.5.4 Olfactory stimulation

Odor delivery was performed with a pressurized superfusion system with Oxycarbon-bubbled (95% O2, 5% CO2, into Oxygen8 perfusion head, Automate Scientific) odor reservoirs, a single odor manifold (Automate Scientific) and valves controlled (ValveLink8.2, Automate Scientific) with a custom written LABVIEW routine. The odorant final concentration at the outlet (4 mm away and pointing toward the nostril ipsilateral to the calcium imaging hemisphere) was always 100 µM. A relatively long stimulus duration was chosen because odors in natural habitats of zebrafish (standing or slow waters in the Indian subcontinent; Spence et al., 2008) are expected to have a long lifetime. We applied odors in 20 s long step pulses, at an inter-trial interval of 3 minutes, while recording calcium activity from pDp or from the lateral pallium (lower zoom acquisition), at a consistent average depth of 50 µm from the lateral surface of the forebrain within the imaging field of view.

### 4.6 Processing of calcium movies

#### 4.6.1 Bidirectional scanning correction

Because scanning mirrors do not follow a linear speed profile, bidirectional scanning can lead to local misalignments between odd and even-numbered lines of pixels, even when the average mismatch is near zero. To correct for this, we used a single reference trial per fish and estimated local shifts based on line-by-line cross-correlation on the time-averaged histogram-equalized (Zuiderveld, 1994) calcium movie from that trial. We then take a consensus inverse transform, which resembled the inverse speed profile of the scanner. Because this mismatch is a property of the scanner and independent of the sample, we could then apply this correction across the subject’s dataset.

#### 4.6.2 Motion artefact correction

We observed that opening a craniotomy tends to compromise the structural rigidity of the skull to shear forces, which may occur during vigorous tail movements under head fixation. Additionally, the exposed brain tissue undergoes rhythmic, often detectable mechanical oscillations (wobble). This is probably due to edding dynamics of the external flow of medium, as well as internal pulsations of the ventricular and vascular systems of the brain. To compensate for these sources of motion, we used a sequential two-stage approach. First, we manually identified all brief time intervals with strong motion artefacts that rendered anatomical information from the target field of view impossible to recover: these intervals are eliminated and replaced with empty image frames. Second, remaining motion artefacts and small rigid image offsets were compensated by applying an optic flow based warping algorithm (Flotho et al., 2022) to align each frame to a time-averaged reference. We computed these transforms on histogram-equalized movies and references (Zuiderveld, 1994) and then applied them to their raw counterparts.

#### 4.6.3 PCA/ICA-based source decomposition (generation of parallel datasets)

The use of small optical components, such as the prism, can reduce the maximum beam cross section and therefore increase laser power demands and limit the effective numerical aperture of the objective lens. These technical limitations can lead to a lower signal to noise ratio and larger point-spread function as compared to direct tissue imaging through a high-NA objective. In practice, this can result in optical mixing effects such as signal contamination of a somatic ROI by out-of-focus neuropil activity. To control for this, we developed a custom singular value decomposition (SVD) / independent component analysis (ICA) based signal source decomposition pipeline, which identifies anatomical modes of temporally correlated signals (Fig. S1f-i). We used this approach, combined with spectral analysis of the resulting independent component loadings maps, to identify anatomically broad components as putative time-varying background or out-of-focus neuropil activity.

We then used only these “background ICs” (bICs) to reconstruct full movies containing only putative background signal, rather than activity from in-focus somata. Subtracting this reconstructed movie from the raw data yielded putatively de-contaminated calcium movies, which were used to extract somatic activity traces. To ensure that the specific choice of bIC subsets did not qualitatively affect our downstream analysis results and conclusions we generated four parallel datasets based on different bIC sets: (1) “raw” (raw data; no source decomposition), (2) “IC1” (bIC set was always defined as the single most time-variant IC, which coincided with the lowest spatial frequency, as expected), (3) “conservative” (a small set of bICs, picked manually on the basis of their spatial profile and variance to contain a minimal amount of signal, putatively minimizing false rejections at the cost of potentially missing some correct rejections), and (4) “liberal” (a more comprehensive set of manually picked bICs, aimed at maximizing correct rejections at the potential cost of false positives). Based on a visual inspection of the resulting movies, we identified the “IC1” dataset as the one with the lowest putative optical contamination and the lowest decrease in fluorescence carrying putative somatic signal. For this reason, we use this dataset for all further analyses. However, other datasets gave similar and the dataset choice did not affect the main conclusions (see Fig. S2 for a comparison of main results obtained using all four procedures).

#### 4.6.4 ÄF/F

Somatic ROIs were defined manually using a custom GUI. Only ROIs that were stable across all trials were included in the analysis. To extract neuronal activity, we first measured and subtracted a global offset given by background input from the PMT. For each ROI, pixel intensities were then averaged for each frame to yield a raw fluorescence trace, F. The calcium activity trace was then defined as ΔF/F = (F-F_0_)/F_0_, with F_0_ indicating the trial-specific 10^th^ percentile of the neuron’s F and indicating its baseline fluorescence intensity (Fig. 1b-c). ΔF/F traces were then deconvolved using CASCADE (Rupprecht et al., 2021) (Fig. 1c) and the resulting inferred firing rates (“Firing rates” in the figures and text, for short) were used for all remaining analyses, unless stated otherwise.

### 4.7 Phenomenological model

To validate the geometric capacity analysis independently of biological variability, we developed a generative model of population neural activity that embeds known geometric structure into a simulated neural manifold. The model produces synthetic calcium imaging recordings of N neurons across K odor stimuli, each presented R times in a defined sequence. We chose this trial sequence to be the same as in our experiment.

#### 4.7.1 Geometric Axes

All geometry is defined in R*^N^*. A *D*-dimensional identity subspace is first established as an [*N* × *D*] orthonormal basis *U*, interpolated between a canonical sparse representation (the first *D* standard basis vectors, yielding single-stimulus neurons) and a Haar-random orthonormal frame (yielding dense mixed selectivity), controlled by a mixing parameter *ρ*_mix_ ∈ [0,1]. Four families of unit vectors are then constructed within or orthogonal to span(*U*):

1. **Identity axes e***_k_* (one per odor, *k* = 1*,…,K*) live inside span(*U*) with prescribed pairwise inner products ⟨**e***_k_,* **e***_j_*⟩ = *ρ_e_* for *k* ≠ *j*.
2. **Novelty axes n***_k_* are constructed as **n***_k_* = **e***_k_* + **w***_k_*, where **w***_k_* ⊥ span(*U*) and ⟨**w***_k_,* **w***_j_*⟩ = *ρ* for *k* ≠ *j*. Because **e***_k_* ∈ span(*U*) and **w***_k_* ⊥ span(*U*), the cross-product ⟨**e***_k_,* **w***_k_*⟩ = 0, guaranteeing ∥**n***_k_*∥ = 1. The parameter *ρ* controls the between-odor novelty-axis correlation.
3. **Baseline axes u***_s_,* **v***_s_* are unit vectors sequentially orthogonalized to the full set {*U,* **n**, **d**} (and to each other) by projecting random vectors onto the complement of the accumulated column space via QR decomposition.

#### 4.7.2 Mean Population Patterns

For trial *t* presenting odor *k* at its *r*-th repetition, the noiseless mean population response vector *µ*(:*, t*) ∈ R*^N^* is:

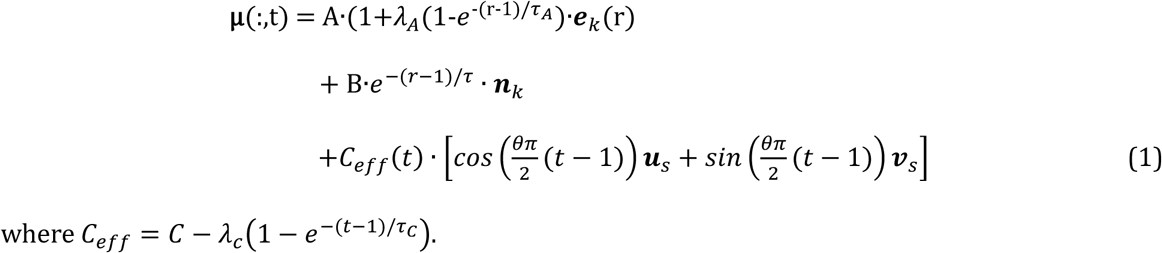

where *C_eff_* = *C* − *λ_c_*(1 − *e*^−(*t*−1)/*ττCC*^).

The *identity* term represents the stable, odor-specific representation, with amplitude *A* that can grow across repetitions (gain *λ_A_*, time constant *τ_A_*).

The *novelty* term captures an initial response of amplitude *B* that decays exponentially with time constant *τ* (in units of repetitions) along an odor-specific axis that may be correlated across odors (via *ρ*). The *baseline* term implements a shared, trial-sequence-wide modulation of amplitude *C*_eff_ that rotates in the plane spanned by (**u***_s_,* **v***_s_*) at angular speed *θπ/*2 radians per trial and can attenuate over the experiment (via *λ_C_* and *τ_C_*). At *θ* = 0 the baseline is a constant global offset; at *θ* = 1 consecutive trial baseline vectors are orthogonal. *µ*(:*, t*) represents the expected firing-rate modulation (in Hz) relative to zero during the odor-response window; a neuron-specific baseline is added by the noise model.

#### 4.7.3 Noise Model

Per-neuron spontaneous firing rates (baselines) are drawn independently from a lognormal distribution with target mean r̄ and coefficient of variation CV_r_. Noise is modelled as a single-factor process combining shared and independent components:

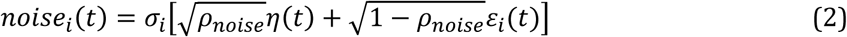

where *η*(*t*) is a shared colored noise process, *ε_i_*(*t*) are independent per-neuron colored noise processes, and *ρ_noise_* is the target noise correlation. The per-neuron noise amplitude *σ_i_* is calibrated to achieve a target Fano factor *F* at baseline. The full synthetic firing-rate trace is assembled as:

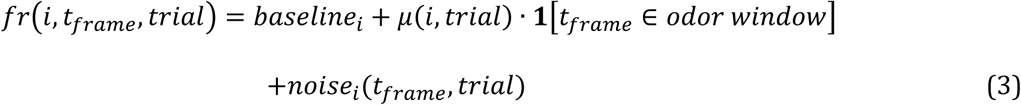

clipped to 0 Hz. Recordings are sampled at 7.67 Hz for 1300 frames per trial, with a 20 s response window, matching the experimental parameters.

#### 4.7.4 Choice of model parameters

A complete set of model parameter values across conditions is available in Table 1.

**Table 1.** Model parameters. Highlighted rows indicate differences across conditions.

| Parameter | Interpretation | Null Model | Naïve-like | Trained-like |
| --- | --- | --- | --- | --- |
| $N$ | Number of simulated neurons | 200 | 200 | 200 |
| $K$ | Number of distinct odors | 6 | 6 | 6 |
| $R$ | Repetitions per odor (basis for the default trial sequence) | 5 | 5 | 5 |
| $D$ | Dimensionality of the shared subspace from which odor-identity axes are drawn | 80 | 80 | 80 |
| $A$ | Amplitude (Hz) of the odor-identity response at the first presentation | 6 | 1 | 1 |
| $\lambda_A$ | Fractional growth of identity-response amplitude with repetition (asymptotic gain = $1 + \lambda_A$) | 0 | 3 | 5 |
| $\tau_A$ | Time constant (reps) governing identity amplitude growth and identity-axis correlation drift | 1.5 | 1.5 | 1.5 |
| $B$ | Amplitude (Hz) of the novelty/habituation response at the first presentation | 0 | 4 | 6 |
| $\tau$ | Decay time constant (reps) of the novelty/habituation response | 1 | 1 | 1 |
| $C$ | Initial amplitude (Hz) of a shared, trial-varying baseline offset | 0 | 5 | 2 |
| $\lambda_c$ | Total attenuation of the baseline amplitude across the session | 0 | 4 | 0 |
| $\tau_c$ | Time constant (trials) of baseline-amplitude attenuation | 10 | 10 | 10 |
| $\theta$ | Angular drift rate of the baseline offset within its 2D rotation plane | 0.06 | 0.06 | 0.06 |
| $\alpha$ | Alignment between each odor's identity and novelty axes (0 = orthogonal) | 0 | 0 | 0 |
| $\rho$ | Pairwise correlation between novelty axes of different odors | 0 | 0 | 0.1 |
| $\rho_e$ | Pairwise correlation between identity axes of different odors | 0.2 | 0.2 | 0.2 |
| $\rho_{\text{mix}}$ | Degree of mixed selectivity (0 = sparse single-odor tuning, 1 = fully dense/random) | 0.7 | 0.7 | 0.7 |
| F | Trial-to-trial variance/mean ratio of single-neuron firing rates | 1.2 | 1.2 | 1.2 |
| $\bar{r}$ | Mean baseline firing rate (Hz) across the population | 0.07 | 0.07 | 0.07 |
| $\text{CV}_r$ | Coefficient of variation of firing rates across neurons (excitability heterogeneity) | 0.9 | 0.9 | 0.9 |
| $\rho_{\text{noise}}$ | Pairwise noise correlation between neurons | 0.1 | 0.1 | 0.1 |
| $\tau_{\text{noise}}$ | Time constant (frames) of temporal smoothing applied to the noise | 8 | 8 | 8 |

All three synthetic datasets simulate a population of 200 neurons responding to 6 odors over 30 trials (5 repetitions each), with response geometry drawn from a shared identity subspace, partial mixed selectivity (*ρ*_mix_ = 0.7), correlated odor-identity axes (*ρ_e_* = 0.2), and a slowly rotating shared baseline component (*θ* = 0.06, *τ_C_* = 10); noise statistics and temporal parameters are identical across conditions.

The three conditions differ in the amplitude and dynamics of odor-evoked responses (Fig. 8a-c). The Null model is obtained by setting *B* = 0, *λ_A_* = 0, and *C* = 0, thereby removing all repetition- and trial-dependent structure from the mean patterns. Under this condition, the population response to any given odor is identical across repetitions: *µ*(:*, t*) = *A* · **e***_k_* for all trials presenting odor *k*. The Naïve-like model has a small identity component that grows moderately with repetition (A = 1, *λ_A_* = 3), a strong novelty-driven component (B = 4), and a large baseline offset that attenuates over the session (C = 5, *λ_C_* = 4). The Trained-like model retains the small identity amplitude but shows the steepest identity growth (*λ_A_* = 5) and strongest habituation (B = 6), a smaller, constant baseline (C = 2, *λ_C_* = 0), and weak positive correlation among novelty axes across odors (*ρ* = 0.1), reflecting training-induced sharpening and stabilization of odor representations. The more pronounced attenuation of the baseline drift component in the Naïve- like compared to the Trained-like model is generally consistent with the experimental observation that odor responses in Naïve animals were more strongly influenced by trial-number-dependent than odor-specific factors.

### 4.8 Analysis

All processing and analysis was performed using a custom pipeline based on MATLAB and Python. Code is available at https://github.com/caudtomm/calcium minimal.

#### 4.8.1 Metrics of activity

We defined single-unit activity metrics as follows.

**Attenuation score** For neuron *i*, the attenuation score *A_i_* is the average across stimuli of the per-stimulus attenuation *A_i,p_*:

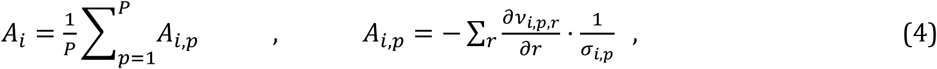

where *v_i,p,r_* is neuron *i*’s average firing rate during the *r*-th repetition of stimulus *p*, and *σ_i,p_* is the SD across repetitions. Negative slopes (decreasing responses) yield positive attenuation scores.

**Selectivity of tuning** Defined as the neuron’s lifetime sparseness *S_i_*:

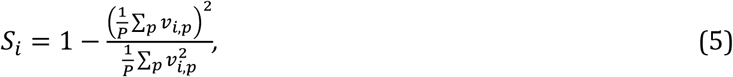

for *P* stimuli. *S_i_* ∈ [0,1], where 0 indicates equal responsiveness to all stimuli and 1 indicates exclusive responsiveness to a single stimulus.

**Stability of tuning** *T_i_* is computed as the average tuning-curve correlation across repetitions. *T_i_* = −1 indicates anticorrelated tuning across repetitions, *T_i_* = 0 randomly varying tuning, and *T_i_* = 1 invariant tuning.

Population activity metrics were defined as follows.

#### Participation ratio

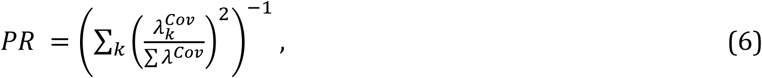

where 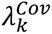 are the eigenvalues of the population covariance matrix during a single-trial odor response.

#### Normalized population sparseness

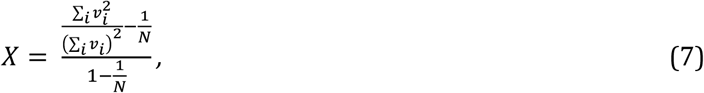

over *N* neurons, as previously described (Vinje and Gallant, 2000; Frank et al., 2019). *X* ∈ [0,1], where 0 indicates that every unit has the same firing rate and 1 indicates that only a single neuron is active.

**Population variance** The variance of *v_i_* across neurons.

#### 4.8.2 Odor classification

Odor identity was decoded from population activity vectors computed as the mean firing rate across the odor-response window for each trial. All classifiers were evaluated using a leave-one-repetition-out cross-validation scheme: for each fold, trials from one repetition index were held out for testing and the remaining trials were used for training. A chance baseline was estimated by repeating each classification 50 times with neuron identities independently permuted within each trial, preserving the marginal activity distribution across neurons while destroying their stimulus-specific structure.

### Template matching (TM)

We computed a class template either as the mean population vector across repetitions for each odor, or as the mean population vector of a single repetition. Test trials were assigned to the most highly correlated template.

### Linear and quadratic discriminant analysis (LDA/QDA)

We fit a Gaussian generative model to each class using *fitcdiscr* (MATLAB Statistics and Machine Learning Toolbox). LDA assumes a single covariance matrix shared across classes, yielding linear decision boundaries between odor classes, while QDA fits a separate covariance matrix per class, allowing curved (quadratic) boundaries at the cost of additional parameters. Test trials were assigned to the class whose fitted Gaussian density was highest. Because the population dimensionality exceeds the number of training trials per class, we used pseudoinverse-regularized covariance matrices (options ’pseudoLinear’/’pseudoQuadratic’), which substitute a Moore-Penrose pseudoinverse for the singular covariance matrix.

### Support vector machine (SVM)

We used *fitcecoc* (MATLAB Statistics and Machine Learning Toolbox) with a linear SVM base learner combined via a one-vs-all error-correcting output code (ECOC) scheme to extend the inherently binary SVM to multiclass odor decoding: one linear classifier was trained per odor to separate it from all others, and test trials were assigned to the class with the highest decision score.

### Direct basis decoder (DBD)

We projected each test trial onto the set of class mean vectors (identical to the TM templates) and assigned the label with the highest projection score.

For all classifiers, decoding performance was quantified as the fraction of correctly classified test trials, averaged across cross-validation folds.

#### 4.8.3 Manifold capacity analysis

Manifold capacity analysis was performed according to the GLUE framework (Chung and Abbott, 2021; Chou et al., 2025) as previously described (Hu et al., 2026), with minor modifications. We first perfomed this analysis separately on manifold datasets constructed according to different definitions (”manifold types”, see Fig. S11a). Next, we optimized two critical sampling parameters for each run: the number of points per subsample (P) and the number of subsampling repetitions (R). Both were determined analytically from the observed manifold sizes in the dataset. Manifold size is defined as the total number of data points available for a given manifold. After excluding frames containing NaN values, which may be introduced during pre-processing (see above), manifold sizes ranged from 400 to 1000 data points per manifold across animals in this study. For each manifold type, we select an appropriate parameter set according to the following rules, taking into account the range of manifold sizes available in the dataset.

### Subsample size (P)

For a given manifold of size M and a dataset minimum number of neurons N_min_, P was chosen as:

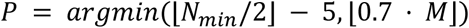

The first term ensures that no single subsample exceeds half the smallest available dimensionality (neuron number; Chou et al., 2025). The second term limits P to 70% of the current manifold size, retaining at least 30% of points as held-out mass. The tighter of the two constraints applies, and the maximum subsample size for odor manifolds was set to 70 data points.

### Repetition count (R)

To determine R, we estimated the minimum number of repetitions required to achieve full manifold coverage as a function of P. Coverage was defined as the fraction of unique manifold points seen at least once across all subsamples. For each combination of P and repetition count, we simulated 100 independent draws of P points sampled uniformly with replacement from a synthetic manifold of size M, and computed the mean fractional coverage. Full coverage was declared when the expected coverage reached or exceeded 99%. This procedure was repeated for repetition counts from 10 to 300 (in steps of 10), tracing the full-coverage frontier F(P) for each manifold size. The operating repetition count was then set to:

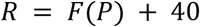

The additive offset of 40 repetitions was included because metric estimation error curves (the variability of the estimated manifold geometry metrics as a function of R) saturated substantially later than the 99% point-coverage frontier. Therefore, oversampling beyond the coverage frontier carries negligible statistical cost while providing a buffer against metric instability.

#### 4.8.4 Dimensionality reduction of neural activity patterns

Before applying PCA or UMAP to our neural data, we z-scored it along the neurons-dimension and removed missing data points. When data was pooled across animals (after subject-specific normalization), missing data from individual fish was replaced with putative inactivity to reduce missing period crosstalk. Additionally, all pooled projections shown were computed and visualized after replacing missing data points with the last-measured firing rate values, which qualitatively did not impact our findings, and were validated at the single-fish level with simple omission of missing data points (not shown). Standard PCA was computed via MATLAB’s *pca* function. For UMAP (Meehan et al., 2025), embeddings were computed using a Euclidean distance metric, min_dist = 0.8, n_neighbors = 199, n_components = 2, and spectral initialization.

## Acknowledgements

We thank SueYeon Chung and Chi-Ning Chou (Harvard University), who generously shared code for GLUE analysis and engaged in insightful discussions. We thank members of the Friedrich lab for fruitful interactions. This work was supported by the Novartis Research Foundation, by the European Research Council (ERC) under the European Union’s Horizon 2020 research and innovation program (grant agreement no. 742576), and by the Swiss National Science Foundation (SNSF; grant no. 310030_212236).

## Supplementary Material

**Figure S1.**
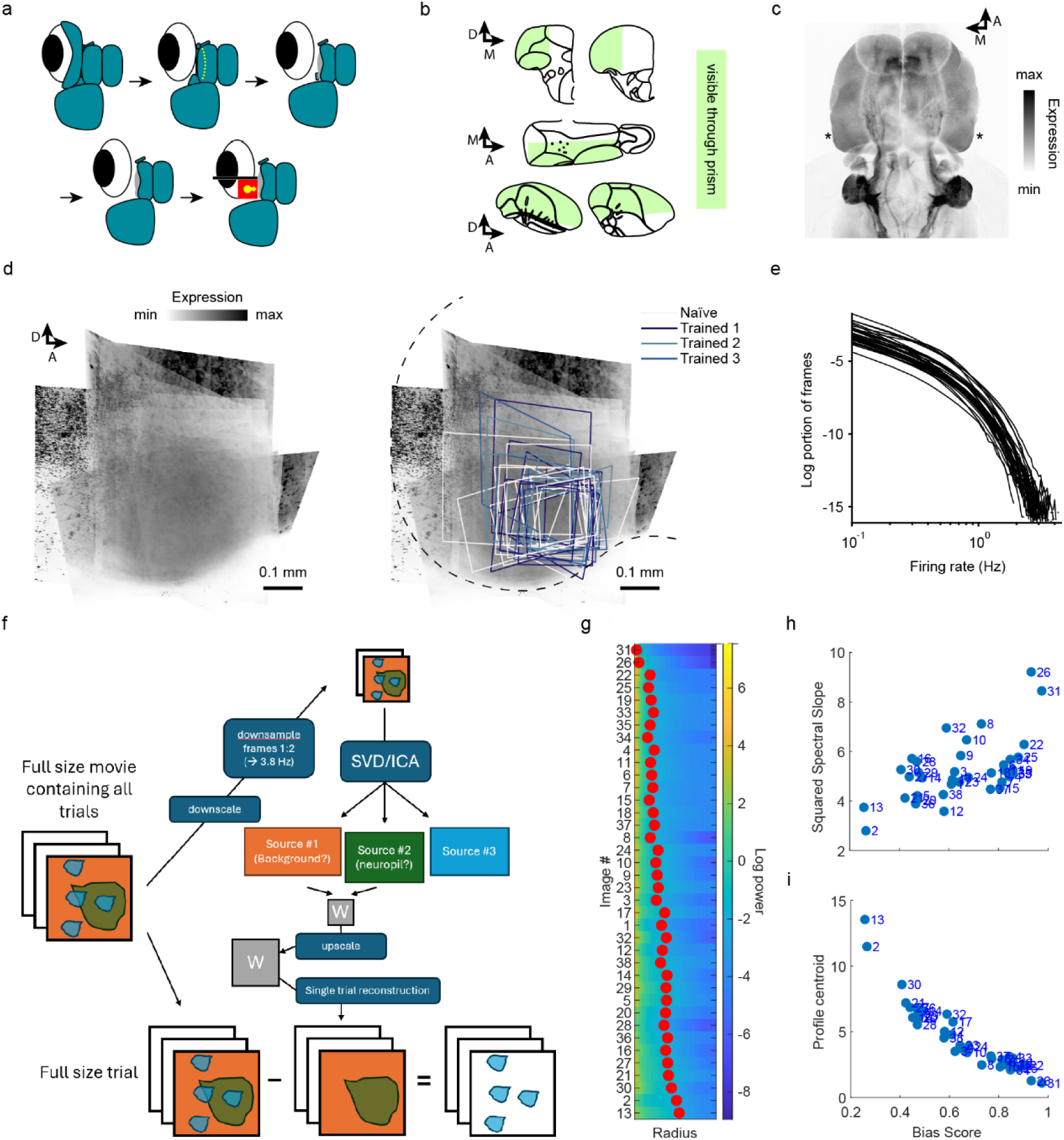
Experimental strategy for physiological recordings. **a** Main steps of surgery and insertion of the prism (red). **b** Schematic outline of optical access to the lateral dorsal forebrain through the prism (shaded areas). Schematics adapted from Wullimann et al., (1996). **c** GCaMP6f expression in the forebrain of an adult Tg[neurod1:GCaMP6f] zebrafish. GCaMP is broadly expressed in the telencephalon (except olfactory bulbs) and habenulae, including pDp (asterisks). Maximum-intensity confocal projection from a full cubic-cleared brain. **d** Maximum intensity projections of low-zoom anatomical 2-photon stacks from all 35 fish (14 Naïve fish and 21 Trained fish; Methods), registered and averaged across fish. Right: Overlay with the co-transformed fields of view of activity measurements. Different outline colors indicate different experimental groups. Dashed line outlines the posterior telencephalic ridge. **e** Inferred firing rate distributions across fish. **f-i** Fluorescence source decomposition. This procedure was used to remove signal components representing putative optical contamination (primarily out of focus fluorescence; Methods) **f** Pipeline based on singular value decomposition and independent component analysis (SVD/ICA). **g-i** Identification of the independent component (image) with the lowest spatial frequency (example of a single fish). This spectral analysis is performed on the spatial loading images for different independent components. **g** Log radial spectral power profiles. This value indicates the average power of the image’s Fourier transform, depending on radial distance from the center of the Fourier-transformed image. As power falls approximately with a logarithm of the radial distance, we extracted the log-centroid (red circles) and slope of this curve (’spectral slope’). Independent components are sorted by decreasing spectral slope. **h** Squared spectral slopes versus spectral bias score. The bias score is computed as P(low freq)/(P(low freq) + P(high freq)), where P is the average power of the 10% lowest or highest spatial frequencies. A high bias score typically reflects a broader spatial profile (consistent with background contamination). **i** Spectral decay centroids (red circles in **g**) versus bias scores.

**Figure S2.**
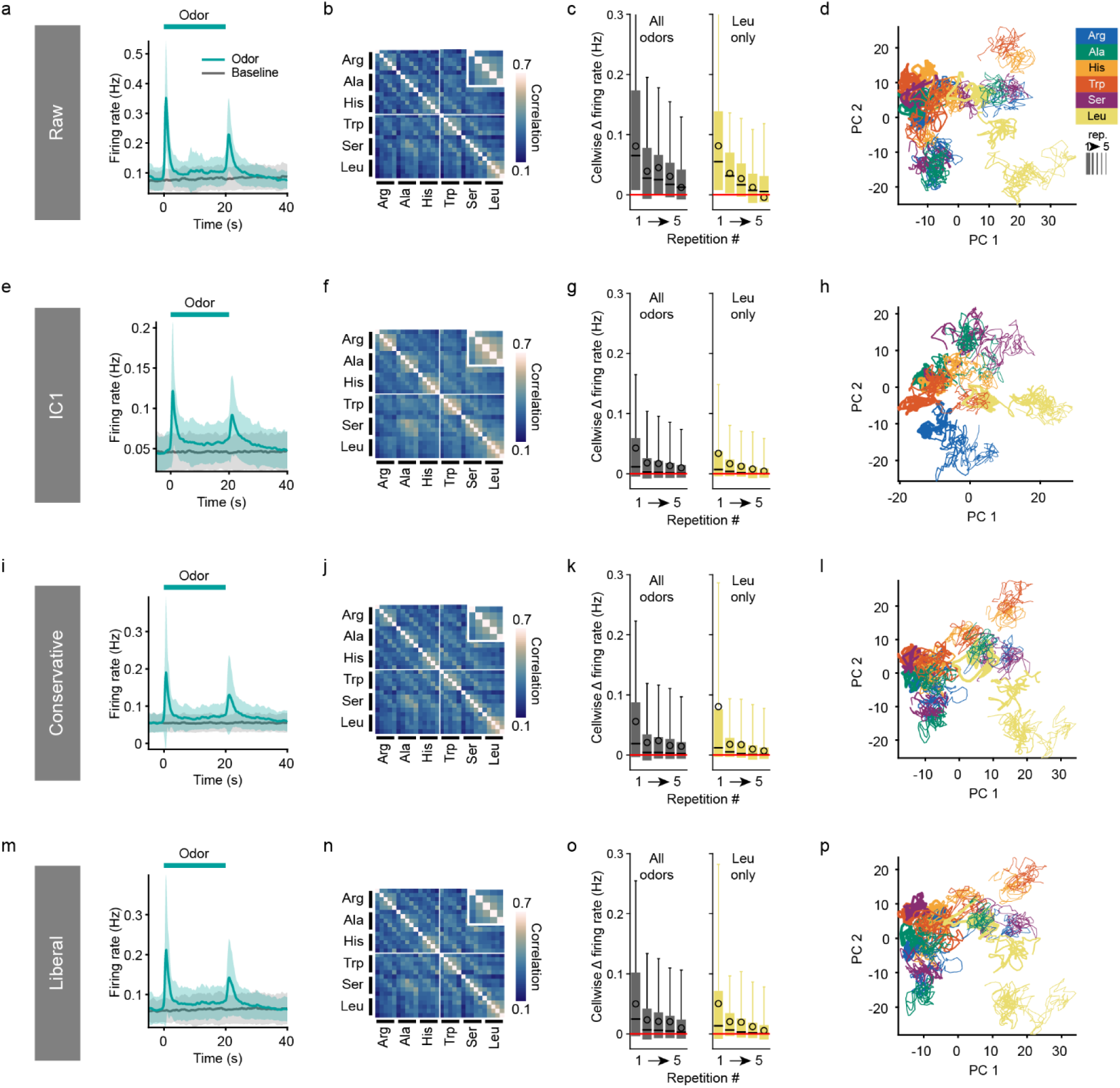
Effects of fluorescence source decomposition parameters on analysis results. **a-d** Key results obtained using raw fluorescence data (all fish combined). **e-h** Same after subtraction of the independent component defined as background “IC1”. **i-l** “Conservative” manually curated set. In each fish ICs were inspected manually, and removed only if no obvious signal could be identified in the spatial pattern. This procedure is expected to minimize false rejections (at the cost of potentially missing some correct rejections). **m-p** “Liberal” manual IC set. In addition to the ICs of the “Conservative” set, this set includes additional ICs, selected manually in each fish to maximize correct rejections (at the cost of potentially including some false rejections). **a,e,i,m** Average single-cell odor response peristimulus time trace (mean ± SD; average over all odors, trials and fish; n = 35 fish including Naïve and Trained groups; Methods). **b,f,j,n** Average inter-trial correlation of time-averaged odor responses. Inset: correlations across same-odor repetitions. **c,g,k,o** Single-cell average baseline-subtracted firing rate during odor responses across repetitions. Left: including all odors. Right: including only responses to Leu (added for the first time at the end of block four; Fig. 1i). **d,h,l,p** PCA embedding of pooled data. Note that subtraction of the independent component representing background (“IC1”) enhances structure in the correlation matrix and the PCA embedding, consistent with removal of non-specific background, but structure is observed also in raw and manually curated (conservative, liberal) datasets.

**Figure S3.**
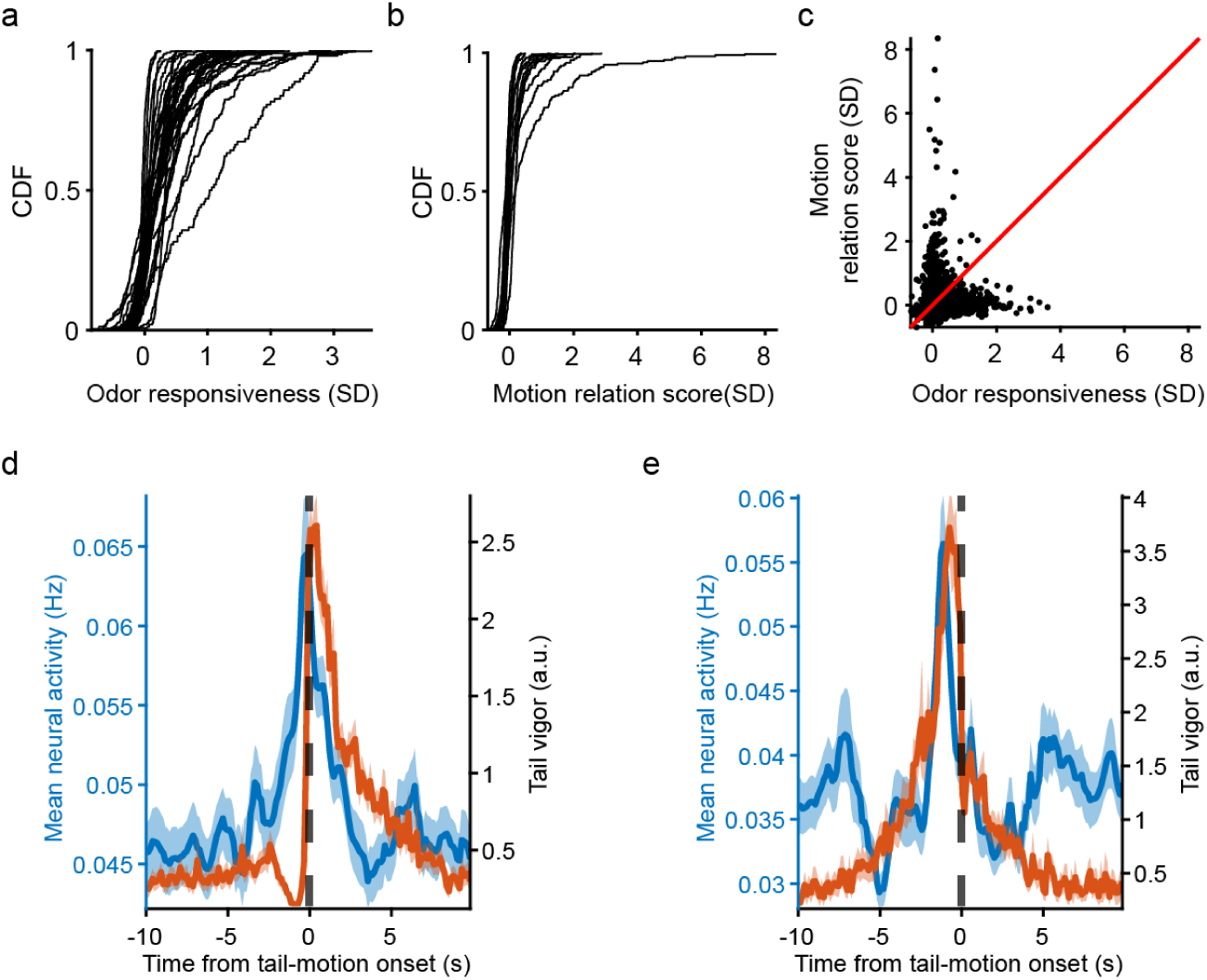
Motor-related activity. **a** Cumulative distribution of cell-wise odor responsiveness scores (average activity during odor responses, z-scored using mean and SD of the full activity, see text). Each curve corresponds to one fish (n = 35 fish including 14 Naïve fish and 21 Trained fish; Methods). **b** Tail motion relation score defined as the average activity during tail motion at least 30 s after odor stimulation offset, z-scored using mean and SD of the full activity (n = 15 fish with behavioral tracking, including 7 Naïve fish and 8 Trained fish; Methods). **c** Odor and motion responsiveness of n = 3123 individual cells pooled across all fish with behavioral tracking (n = 15). Linear mixed effects model regression: slope = 0.055, Pearson correlation coefficient = 0.042, p = 0.024. Red line: equal relation to motion and odors. Note that both odor- and motor-related activity was observed but the correlation was low, indicating that odor- and motion-related activity are largely independent. **d** Average inferred firing rate in pDp, triggered upon swim bout onsets in the absence of odor stimulation (at least 30 s after odor stimulation offset; n = 287 onsets, mean ± s.e.m.). **e** Same as **d**, but triggered on the offset of swim bouts (n = 110 offsets). Together, these results indicate that motor-related modulation of activity coincides with odor presentation but affects a component of population activity that is largely unrelated to odor responses.

**Figure S4.**
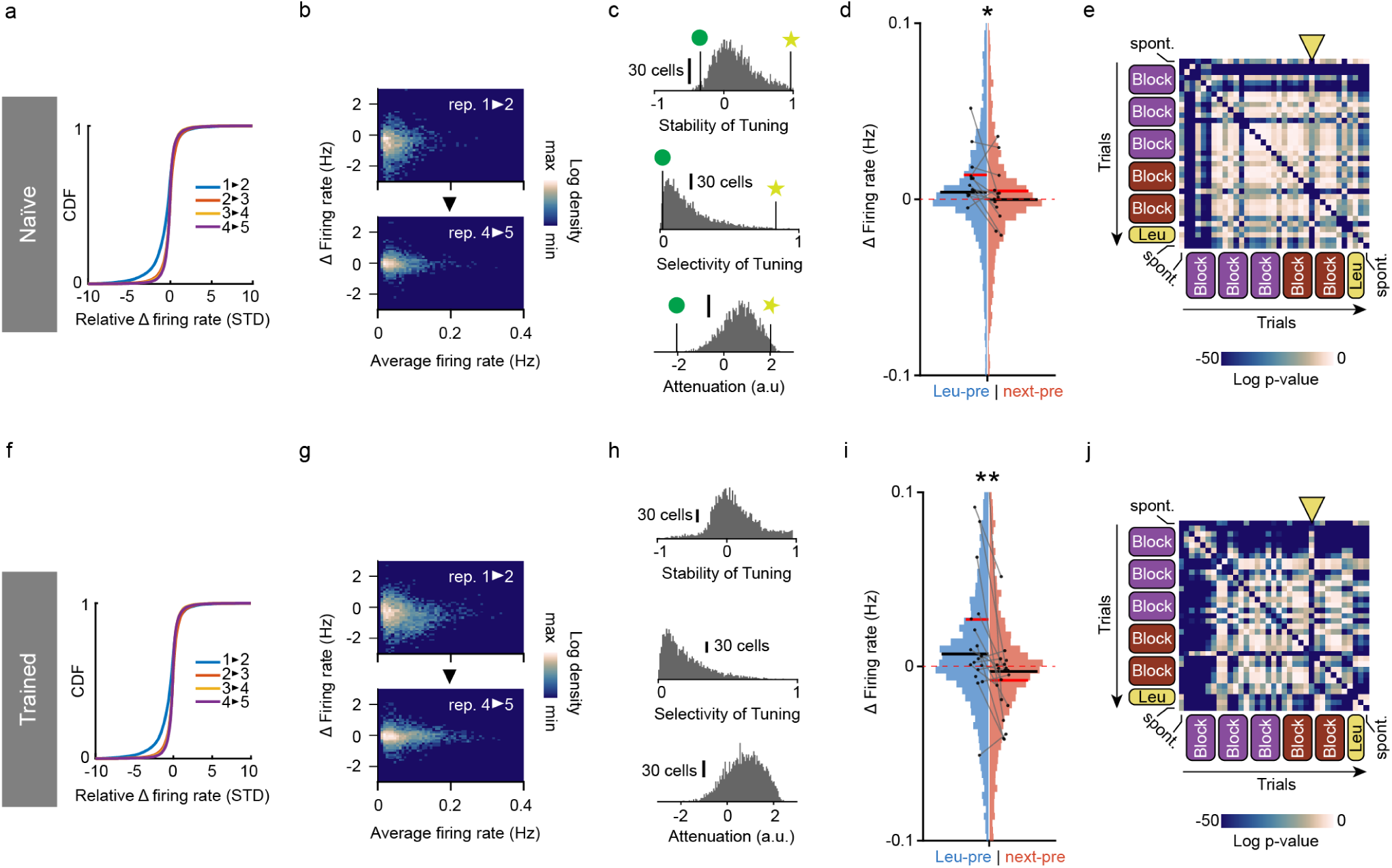
Attenuation and tuning of single-cell odor responses. **a-e** Naïve fish. **f-j** Trained fish. **a,f** Cumulative distributions of relative changes in single unit activity between successive repetitions of the same odors. Negative firing rate changes were more prominent between repetitions 1 and 2 than between subsequent repetitions. **b,g** Single-cell firing rate change after the first (top) and fourth (bottom) repetition of the same odor as a function of the cell’s average activity during odor presentation. Negative changes in firing rate were more prominent after the first repetition with a minor dependence on the average activity (**b**: Pearson correlation, trial 1→2: r = 0.102, p = 3.51·10^-8^; trial 4→5: r = -0.036, p = 0.053. **e**: Pearson correlation, trial 1→2: r = -0.056, p = 1·10^-4^. On trial 4→5: r = -0.007, p = 0.611). **c,h** Histograms of single-cell tuning and attenuation scores (Methods). Green circles and yellow stars indicate the low and high-scoring example cells in Fig. 1j. **d,i** Higher activity upon the first presentation of Leu, compared to neighboring trials. Difference in odor-evoked firing rate change between trial #25 (first repetition of Leu) and the previous trial #24 (“Leu-pre”, blue), or between trial #26 and trial #24 (control, “next-pre”, red). Pooled data from all Naïve (**d,** n = 2892 cells) or Trained (**i,** n = 4635 cells) fish, clipped between ±0.1 Hz for visualization. Red horizontal lines: mean; black: median; connected scatter: single-fish means across cells. 2-sided Wilcoxon signed rank test across single fish averages: p(Naïve) = 0.047, p(Trained) = 8.6·10^-3^. **e,j** Pairwise ΔFiring rate comparisons of pooled data across trials in Naïve (**e**) or Trained (**j**) fish. Log p-values of FDR-corrected pairwise Mann-Whitney U tests, clipped at log(p) = -50 for visualization. Trials are sorted chronologically; yellow triangle indicates the first Leu trial. Note that differences in response intensities are maximally significant with respect to block 1 trials or to the first repetition of Leu, particularly in Trained fish. Taken together, these results indicate that a prominent response attenuation takes place from the first to subsequent repetitions of each odor, and that this attenuation is not constrained to a subpopulation of cells but distributed over the pDp network, independently of average activity or odor-identity tuning.

**Figure S5.**
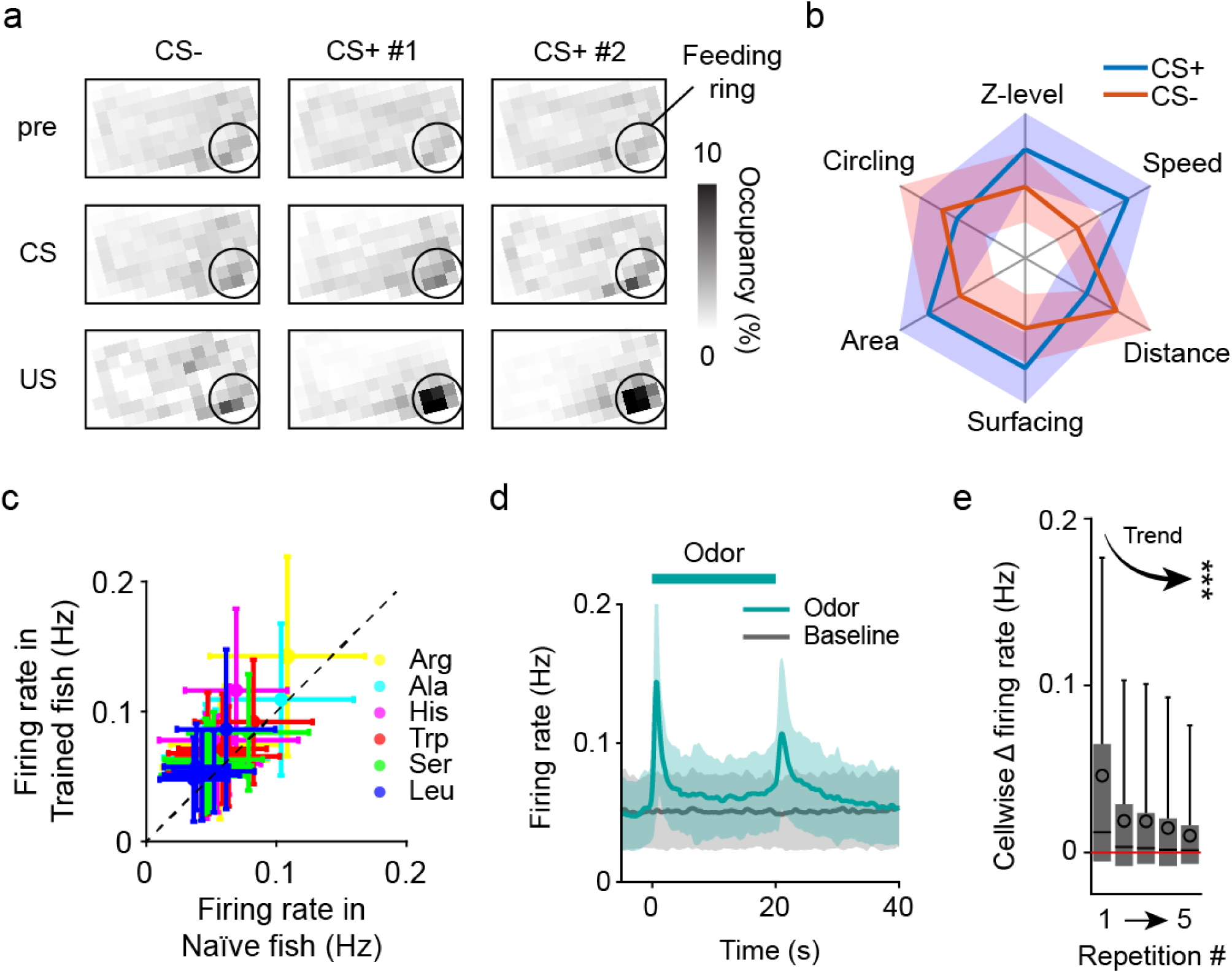
Discrimination training and basic analysis of odor responses. **a** Relative occupancy maps of a trained fish before (pre; 2 min), during (30 s) or after odor presentation (individual training). Note a stronger preference for locations near the feeding ring during presentation of CS+ than CS-. A strong preference for locations near the feeding port is evident during presentation of the US (food; following CS+). **b** Multiday distribution of each behavioral metric associated with appetitive behavior (distance to the reward location, swimming speed, distance to the odor delivery tube, upwards translocation within the water column, surface sampling, circling in the tank) in experienced animals during exposure to the CS- and CS+. Appetitive behavior is indicated by decreased circling and distance from the odor port (“Distance”) and increases in the other metrics. **c** Single-cell firing rates during odor responses in Trained vs Naïve fish per trail. Mean± s.e.m. Red dotted line: equal firing rates. **d** Average single-cell odor response peri-stimulus time trace across Trained fish (n = 21 single-fish trial-averaged traces, mean ± SD) **e** Average single-stimulus firing rate change across repetitions, across all odors in Naïve fish. Friedman test: p < 10^-10^.

**Figure S6.**
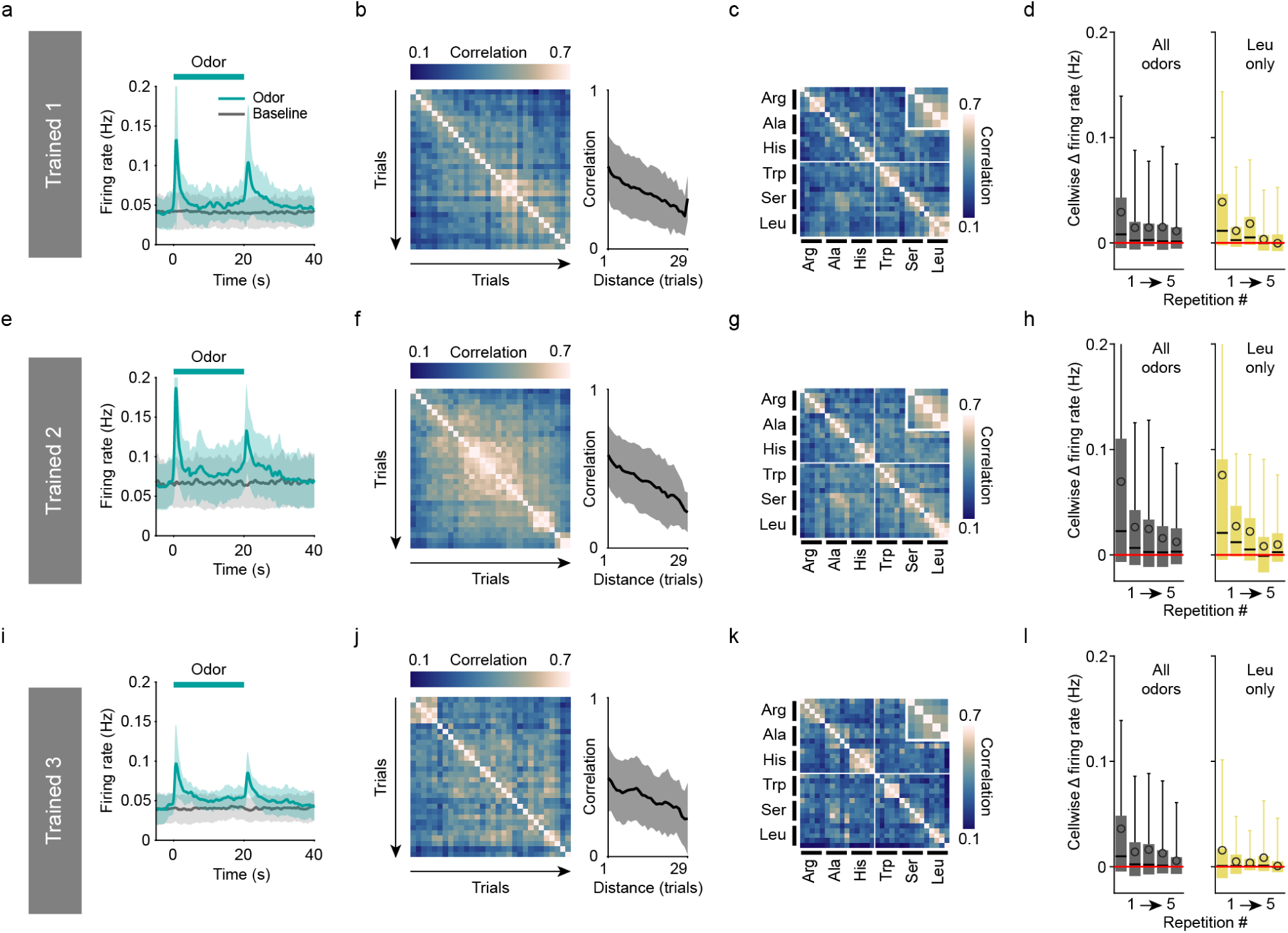
Summary of key data visualizations across training groups. **a-d** Summary plots for training group “Trained1.” **e-h** Group “Trained2”. **i-l** Group “Trained3”. **a,e,i** Average single-cell odor response peri-stimulus time trace (n = 8 Trained1 fish, n = 8 Trained2 fish, n = 5 Trained3 fish, mean ± SD). **b,f,j** Left: inter-trial correlation between time-averaged pre-stimulus activity vectors. Trials are sorted chronologically. Right: inter-trial correlation as a function of temporal distance (mean ± SD). **c,g,k** Average inter-trial correlation of time-averaged odor responses, sorted by odors. Inset: correlations across same odor repetitions. **d,h,l** Single-cell average baseline-subtracted firing rate during odor responses across repetitions, clipped to 0.2 Hz for visualization. Left: including all odors. Right: only including responses to Leu.

**Figure S7.**
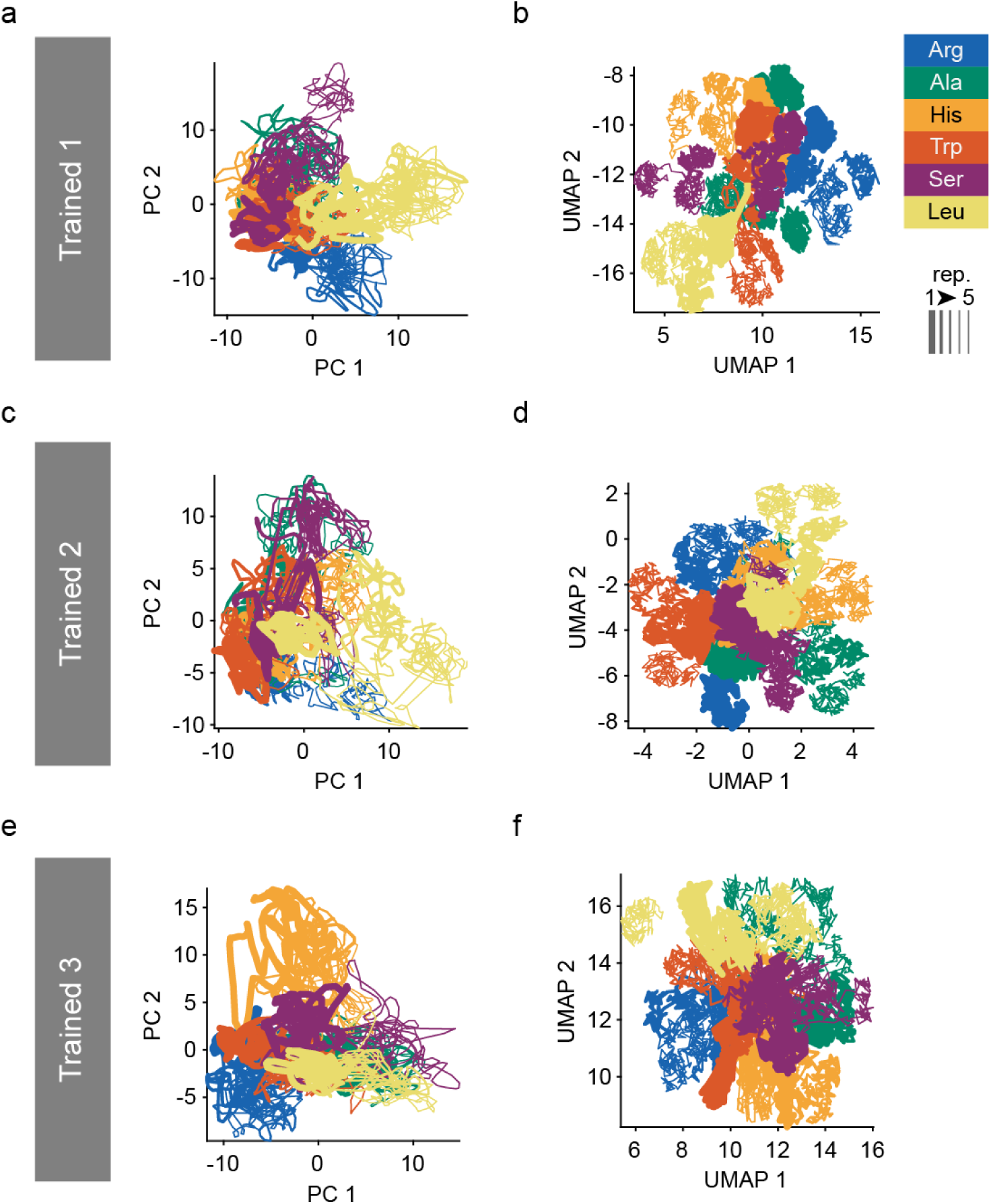
Low-dimensional embedding of pooled response data across training groups. **a-b** Group ‘Trained1” **c-d** Group “Trained2”. **e-f** Group “Trained3”. **a,c,e** PCA embedding. **b,d,f** UMAP embedding.

**Figure S8.**
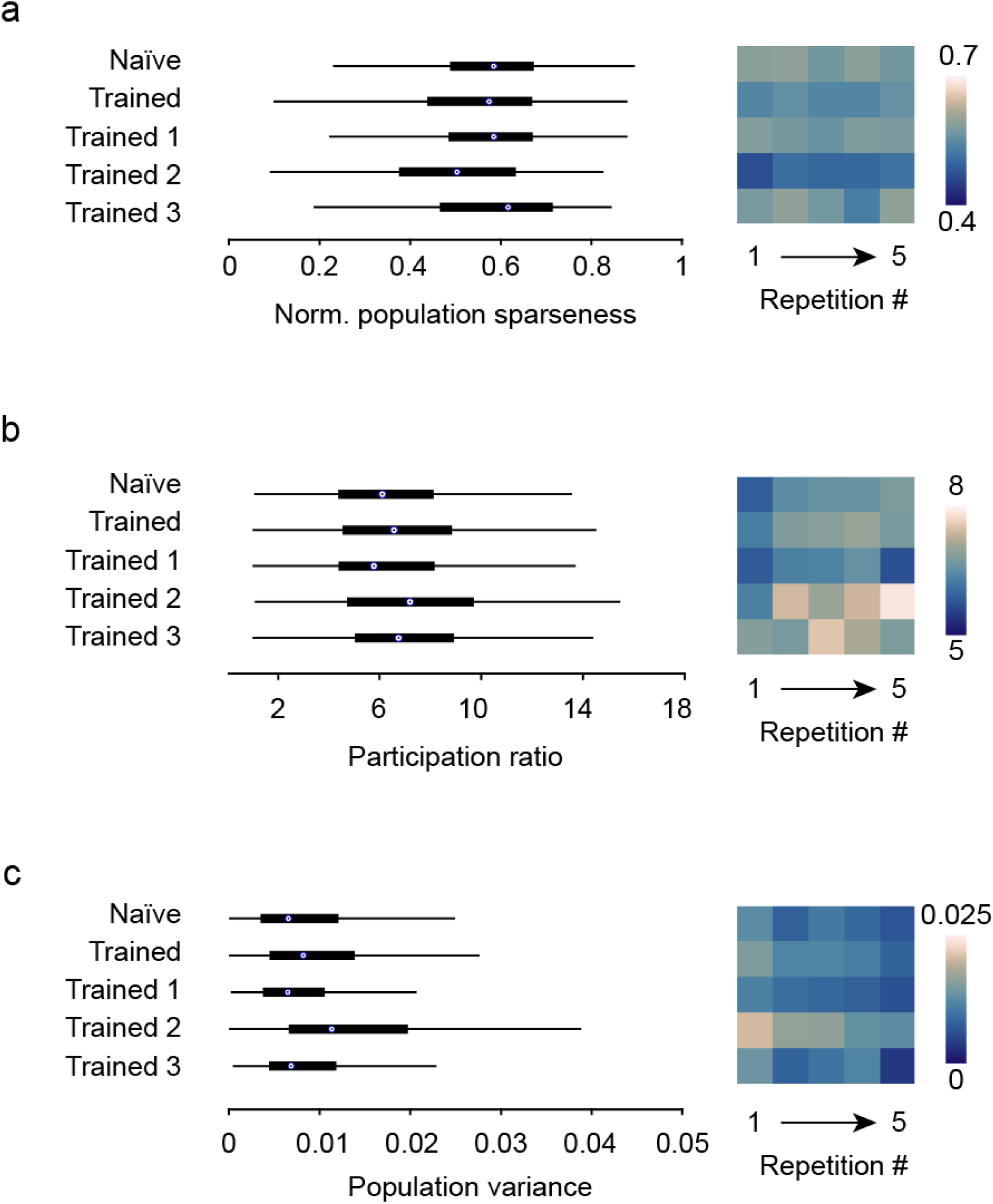
Additional population activity metrics. **a** Normalized population sparseness. **b** Participation ratio. **c** Population variance. Each metric was computed using the time-averaged odor response vector from each trial (see Methods). Left: combined distribution of all trials across training groups. Right: averages across repetitions.

**Figure S9.**
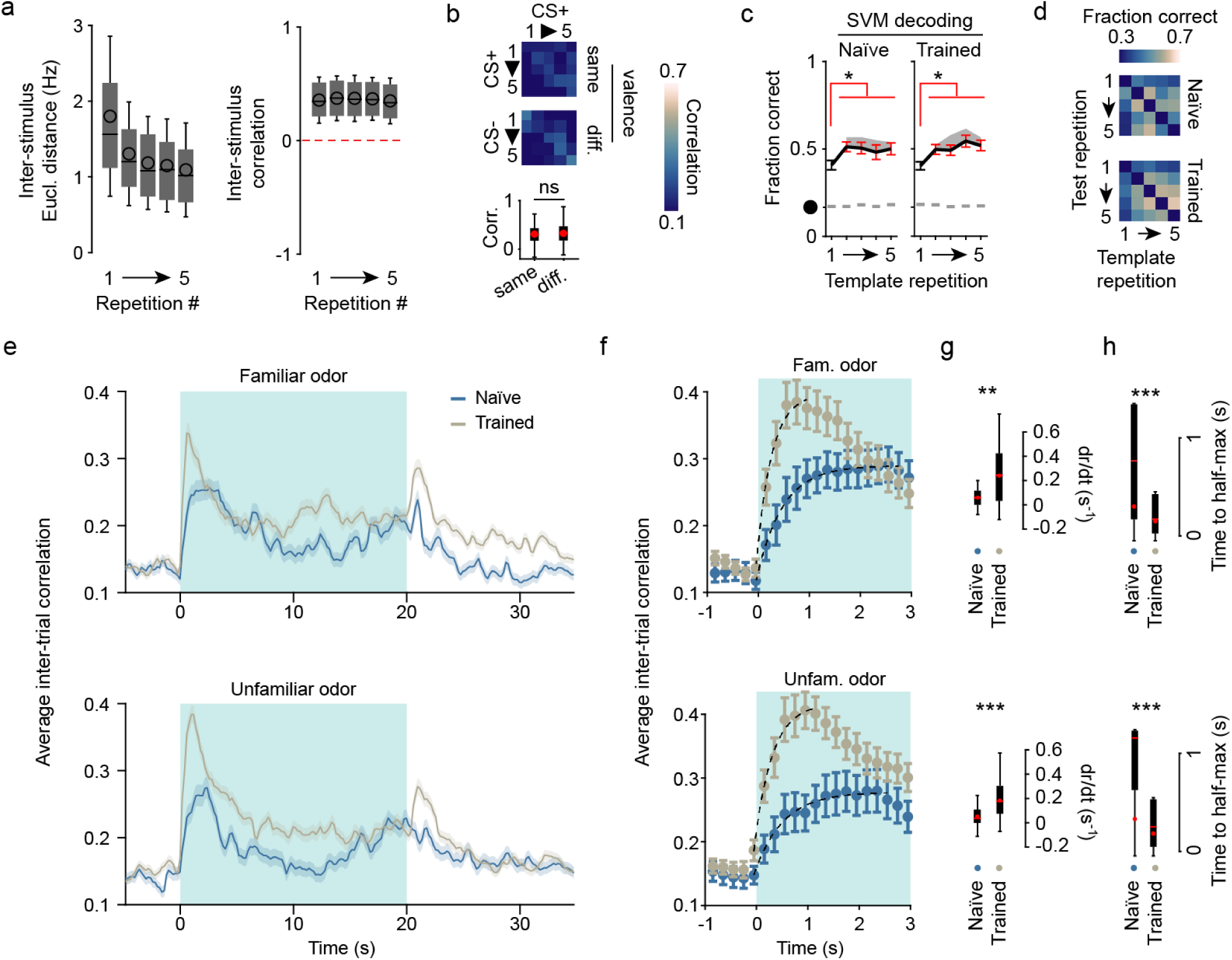
Similarity and decodability of odors across training groups. **a** Similarity of time-averaged response vectors of different odors across repetitions. Left: Euclidean distance decreased over repetitions (from 1.80 Hz to 1.09 Hz over 5 repetitions, on average. Friedman test: p = 1.69·10^-51^), consistent with attenuation of responses. Right: correlations were approximately static (0.36±0.20 mean±SD, Friedman test: p = 0.088). Black lines: median; empty circles: mean; error bars: mean±SD. **b** Pearson correlations across trials of different familiar odors with the same or different valence (CS+ vs CS+ and CS+ vs CS-, respectively), independent of identity. Bottom: 2-sided 2-sample Student t-test (p = 0.297). **c** Fraction of correct classifications in Naïve (left) and Trained (right) fish, obtained by training support vector machines on single trial response patterns (x-axis) and testing on all remaining repetitions of each odor (mean ± s.e.m.; n = 14 Naïve fish or 21 Trained fish; paired Wilcoxon signed rank tests between performance using repetition 1 or another repetition as template: red error bars indicate p<0.05). Black circle: chance level; gray dashes: average performance of 50 equivalent decoders using shuffled unit IDs; gray shaded area: performance increase when excluding repetition 1 trials. **d** Fraction of correct template matching classifications in Naïve (top) and Trained (bottom) fish as a function of template repetition and test repetition. Values are lower on both the first row and the first column, indicating that the first repetition pattern is less efficient for odor identification than subsequent repetitions both as a template and as a target. **e** Inter-trial correlation between activity patterns as a function of time for familiar (top) and unfamiliar odors (bottom) in Naïve and Trained fish. **f** Onset dynamics of intertrial correlations between trials of the same familiar (top) or unfamiliar (bottom) odor. Dashed line: exponential fit to the average curve. **g** Distribution of slopes from single trial-pair onset-to-peak line fits, across groups (Mann-Whitney U test). **h** Time to half-maximum intertrial correlation, estimated by fitting an exponential to the intertrial correlation curve between the odor onset and correlation peak (Mann-Whitney U test). Together, **f-h** confirm that correlational dynamics is faster after training, and that this effect generalizes to unfamiliar odorants.

**Figure S10.**
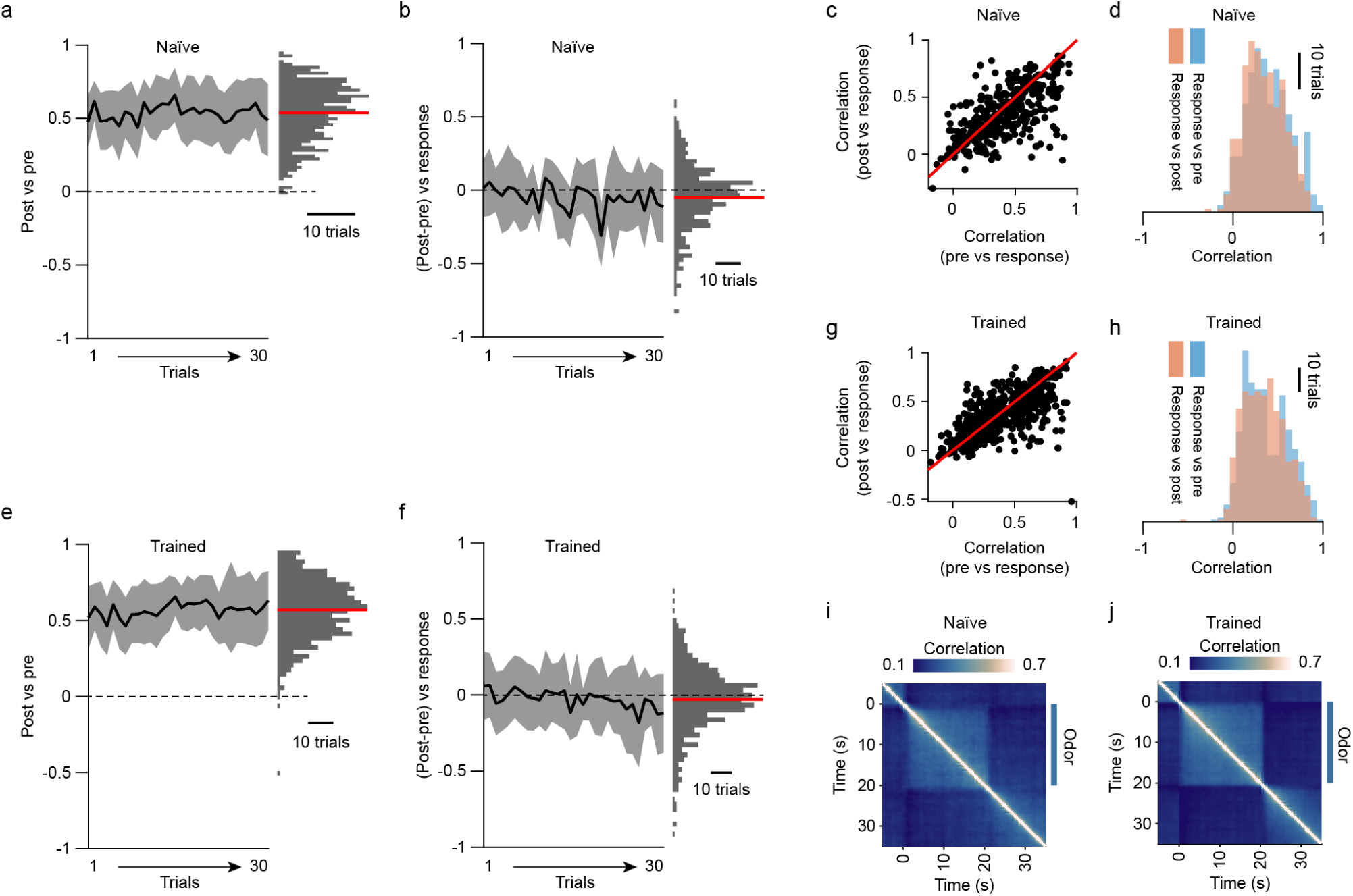
Temporal structure of individual odor responses. **a** Correlation across time-averaged activity from a 20 s pre-stimulus and a 20 s post-stimulus window (starting 65 s after odor offset) in Naïve fish. Pre and post-stimulus activity were correlated (r = 0.54±0.20), with no clear trend across trials. The magnitude of this correlation was similar to the correlation between pre-stimulus activity in consecutive trials (Fig, 6c,d). **b** Correlation between time-averaged stimulus-response activity and difference (post-pre) activity. These activity patterns are uncorrelated on average (-0.05±0.22) with no clear trend over trials, indicating that responses to stimuli do not directly impact subsequent baseline activity. **c** Correlations between stimulus responses and their respective pre and post baseline. **d** Histograms of the individual distributions in **c**. Responses were slightly more correlated to baseline activity before rather than after stimulus application, further indicating that baseline activity did not directly reflect previous odor responses. **e-h** Same as **a-d**, but in Trained fish. **e** Post vs pre correlations: r = 0.57±0.19. **f** (Post-pre) vs stimulus correlations: r = -0.03±0.22. **i-j** Single-trial population activity correlations as a function of time around the odor pulse, averaged over all odor responses. **i**: Naïve fish (n = 420 responses). **j**: Trained fish (n = 630 responses).

**Figure S11.**
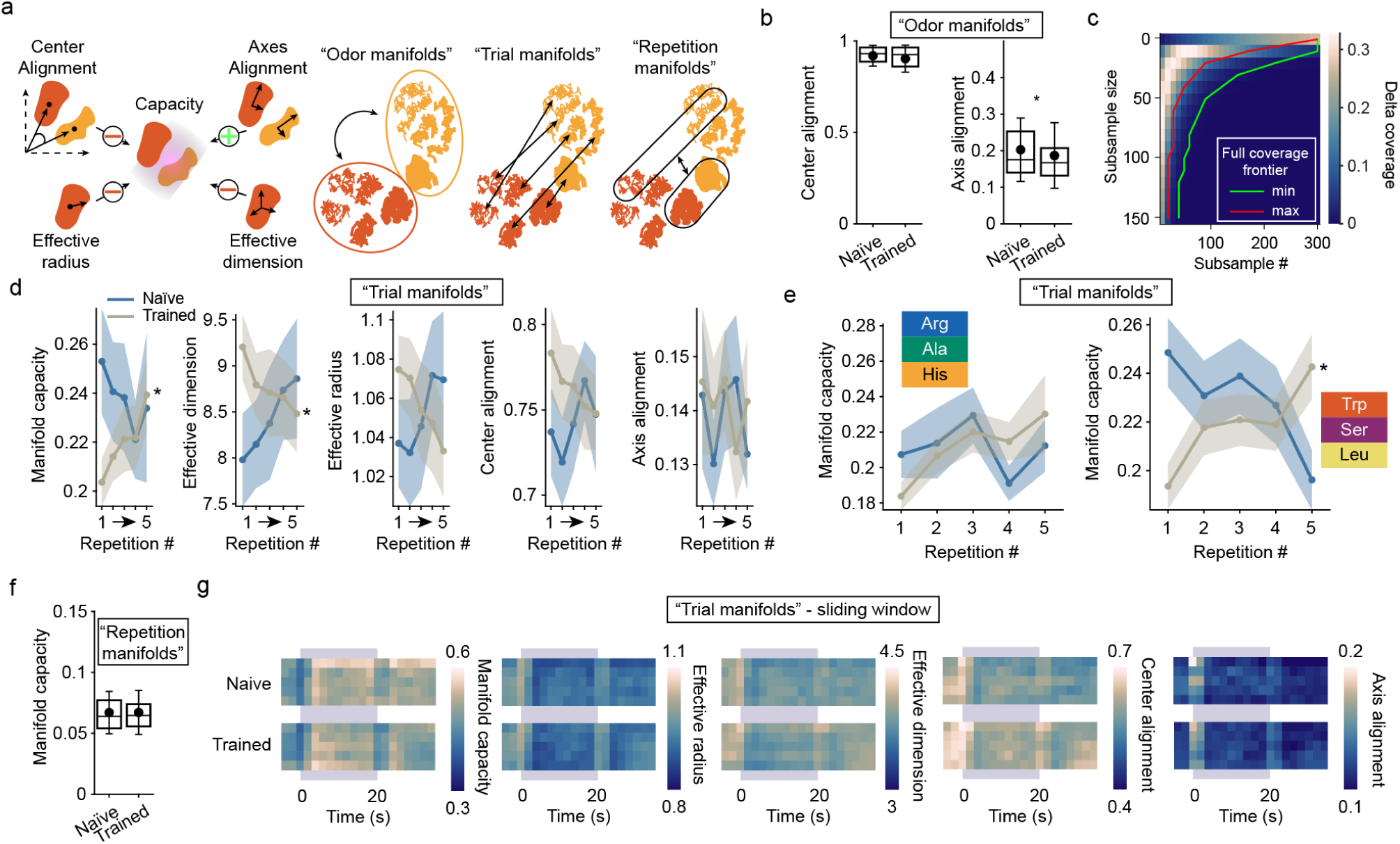
Manifold capacity analysis by GLUE theory. **a** Left: schematic of manifold geometry metrics and their influence on manifold capacity. See (Chou et al., 2025) for details. Right: Manifold construction and nomenclature. “Odor manifolds”: each manifold is the point cloud consisting of activity vectors of all frames during presentation of the same odor, pooled over trials (six odor manifolds per fish). “Trial manifolds”: each manifold is the point cloud corresponding to a single response in one fish (30 odor manifolds per fish). Trial manifold geometry was always measured by comparing same-repetition manifolds. Sliding time window analyses were performed on “sub-trial” manifolds (temporal bins) and compared for the same bins across trials. “Repetition manifolds”: similar to odor manifolds but the grouping variable was repetition number, not odor identity. **b** Center and Axis alignment of odor manifolds across groups (Mann-Whitney U test, p = 0.070 and p = 0.020 respectively, extends Fig. 3m). **c** Expected relative coverage of activity manifolds, based on the sub-sampling strategy (Methods). Heatmap represents the difference between expected coverage of manifolds on opposite extrema of the size spectrum for this dataset (400-1000 available data points per manifold). The lines indicate the full coverage frontier for manifolds of different sizes (minimum and maximum size for this dataset). **d** Quantification of manifold capacity and geometry across repetitions. Friedman test for each curve (mean ± SD): Capacity, p(Naïve) = 0.397, p(Trained) = 0.042; Dimension, p(Naïve) = 0.493, p(Trained) = 0.044; Radius, p(Naïve) = 0.473, p(Trained) = 0.166; Center alignment: p(Naïve) = 0.938, p(Trained) = 0.395; Axis alignment: p(Naïve) = 0.711, p(Trained) = 0.295. **e** Manifold capacity, same as in **d**, left, but broken down by stimulus groups. Left, familiar (Friedman test: p(Naïve) = 0.627, p(Trained) = 0.730). Right, unfamiliar (Friedman test: p(Naïve) = 0.161, p(Trained) = 0.035). **f** Manifold capacity of repetition manifolds across groups. Mann-Whitney U test, p = 0.971). **g** Quantification of manifold capacity and geometry of sub-trial manifolds using a sliding time window. Trial manifolds are defined as in **a,d,e**; each sub-manifold is defined as the section of the trial manifold point cloud contained within a specific time bin.

**Figure S12.**
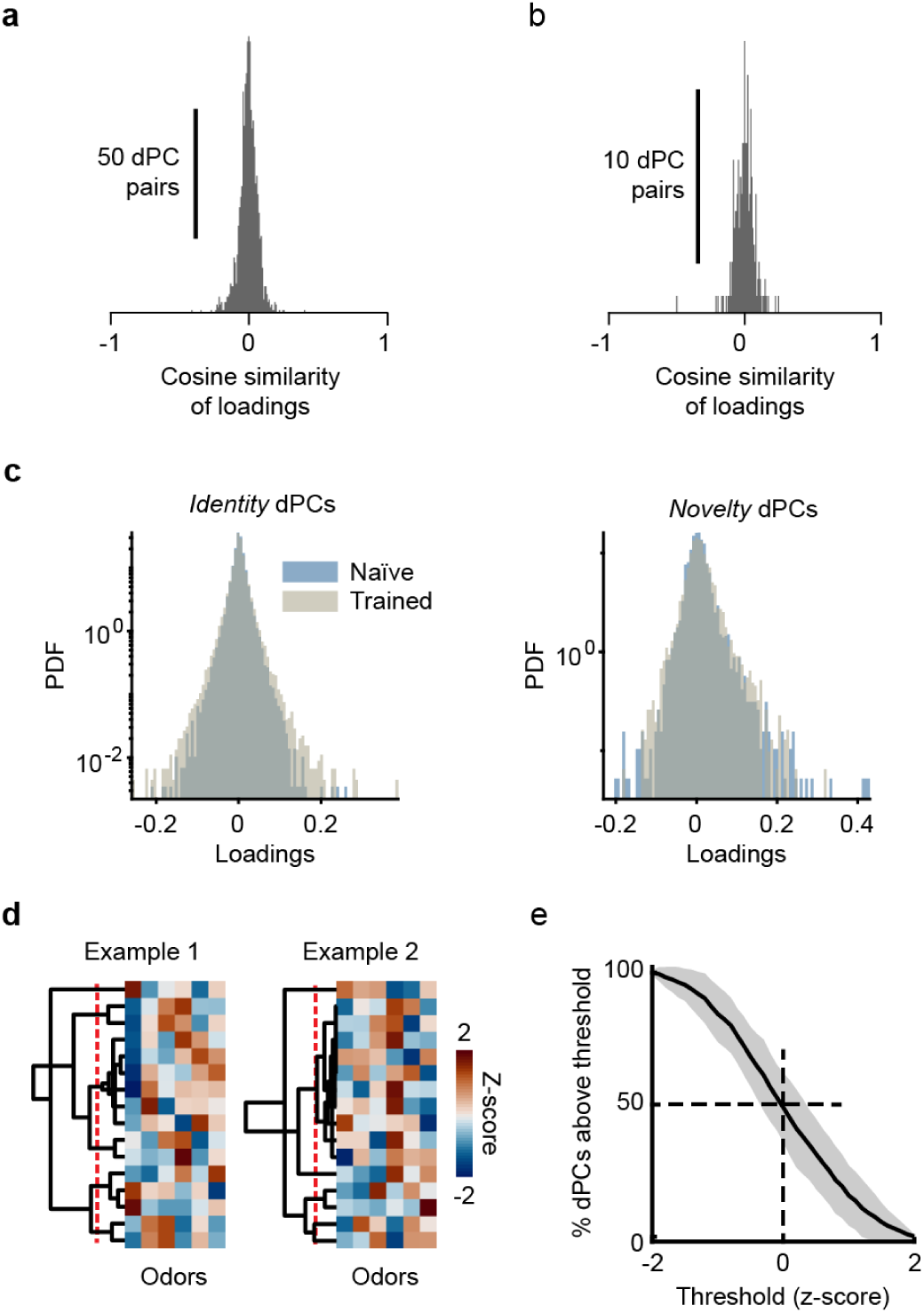
Demixed principal component analysis (dPCA): additional observations. **a** Cosine similarity of reconstruction loadings across *identity* dPC pairs in Naïve fish. As values are close to zero, the reconstruction modes are close to orthogonal. **b** Same as **a**, but for *novelty-identity* dPC pairs, showing that reconstruction modes for *novelty* dPCs are near-orthogonal to the same fish’s *identity* dPCs. **c** Distribution of *identity* (left) or *novelty* (right) dPC loadings across Naïve and Trained fish (semi-log scale). **d** Tuning of *identity* dPCs to different odors in two different fish (representative examples). The activation of each dPC by each odor is represented by z-scores . Note that each stimulus is represented by a combinatorial activation pattern of multiple dPCs. dPCs are ordered by hierarchical clustering based on the cosine similarities of their tuning curves (left). Dashed red-line: cluster cutoff for the formation of 6 clusters (same as number of stimuli). **e** Percentage, for each stimulus, of *identity* dPCs activated above a sliding threshold (mean ± SD). The threshold goes from a low to a high relative value, which are exceeded by most or no dPCs, respectively. Therefore, the curve connecting these extrema provides information on how clustered the *identity* dPC is, relative to odor identity. The broad near-linear section of this curve indicates a distributed representation of odor identity across identity dPCs (a sparse representation would be reflected by step-like transitions).

**Figure S13.**
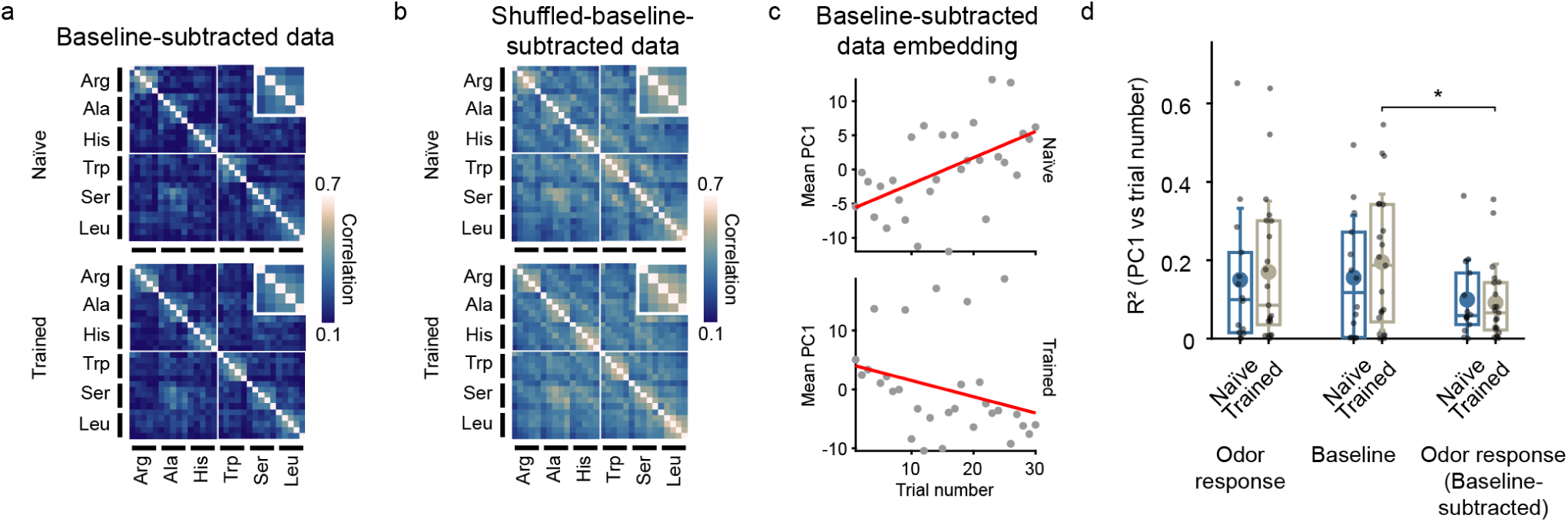
Baseline activity contributes to odor responses and accounts for representational drift. **a** Average inter-trial Pearson correlations between activity patterns during stimulus presentation after subtraction of the baseline activity pattern from each trial in Naïve (top) and Trained (bottom) fish. The Naïve heatmap is the same as in Fig. 6e, inset. Insets: correlations between same-odor repetitions, averaged over odors. **b** Same as **a**, but after subtraction of cell-shuffled baseline patterns. The Naïve heatmap is the same as in Fig. 6f, inset. **c** Mean PC1 scores of pooled baseline-subtracted data from Naïve (top) and Trained (bottom) fish as a function of trial number (see Fig. S15b and Fig. 6g-h for comparison). Red line: linear regression. Pearson correlation values: r(Naïve) = 0.532, p(Naïve) = 2.47·10^-3^; r(Trained) = -0.296, p(Trained) = 0.112. **d** Relationship between PC1 and trial number in individual fish. Because the value of the slope is arbitrary, a comparison of slopes across fish is not meaningful. We therefore compared the squared correlation (R^2^ values). Mean R^2^: full-data odor responses, Naïve = 0.150, Trained = 0.170; pre-stimulus baseline, Naïve = 0.156, Trained = 0.195; baseline-subtracted odor responses, Naïve = 0.099, Trained = 0.091. FDR-corrected Mann-Whitney U test, *: p*<*0.5.

**Figure S14.**
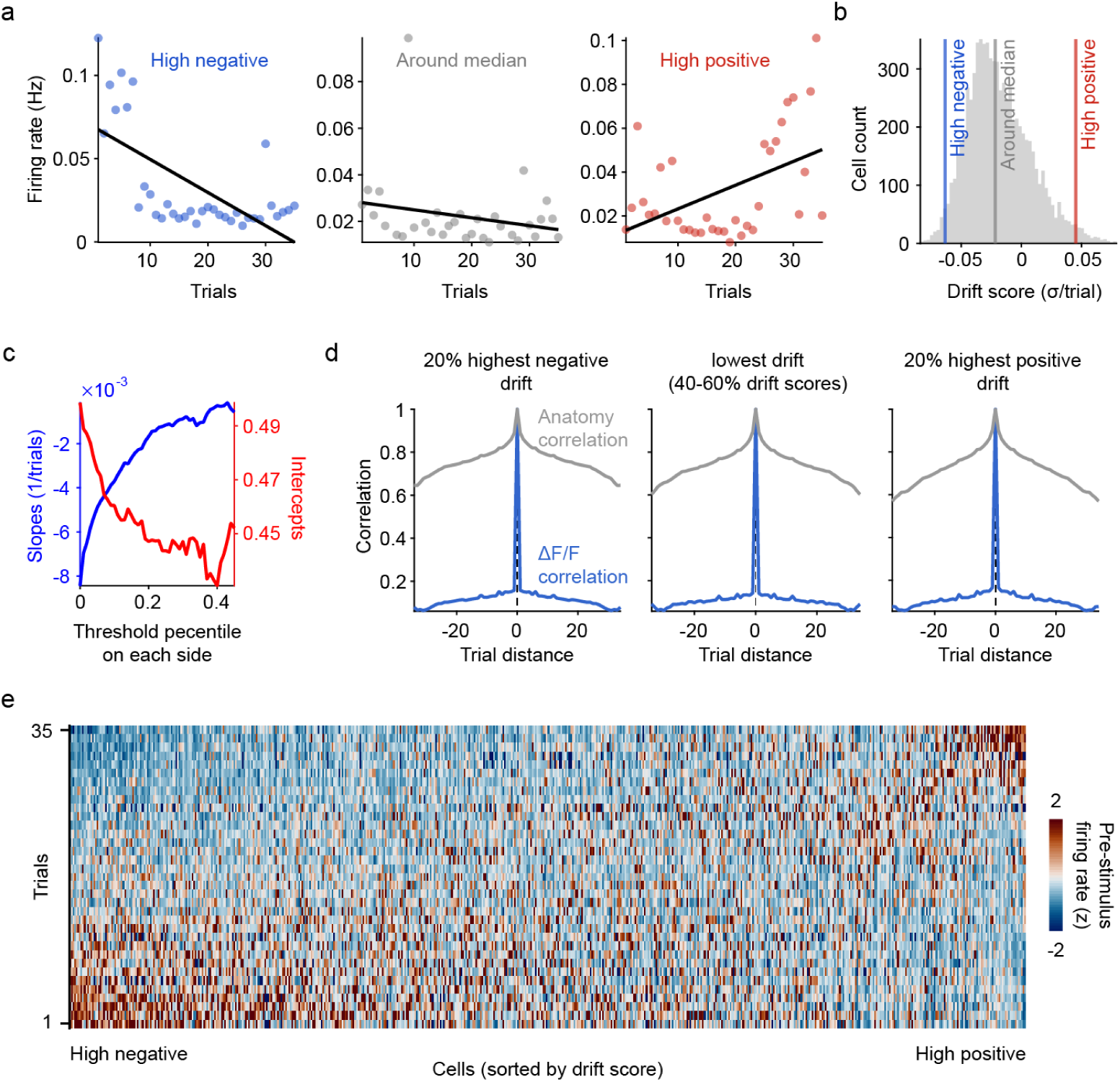
Contributions of individual neurons to baseline drift. To assess whether drift of population activity in the absence of odor stimulation (baseline drift) is caused by a small number of neurons (outliers) we defined a drift score for each neuron. This score is defined by the vectorized ordinary least squares slope of the mean baseline rate (20 s) as a function of trial number, normalized by the SD of average firing rates over trials. **a** Examples of neurons with a high negative drift score (left; marked decrease in baseline firing rate over trials), a neuron with a drift score near zero (center; no major change in baseline activity), and a neuron with a high positive drift score (right; increase in baseline firing rate over trials). **b** Distribution of drift scores, pooled over all fish. **c** To determine how baseline activity patterns and baseline drift depend on neurons with high positive or negative drift scores we removed neurons ranked by their drift score. We then assessed effects on the baseline correlation as a function of trial number (see Fig. 6c,d) by linear fits. Excluding an increasing fraction of neurons with high positive or negative drift scores gradually decreased the correlation between successive baseline patterns (intercept) and the rate at which patterns changed (slope) until approximately 50% of the neurons (25% highest and 25% lowest drift scores) were removed. However, even after removing 50% of the neurons, correlations between successive baseline patterns remained high (∼0.45; intercept). Moreover, the gradual decrease in drift rates (slope) shows that drift was not dominated by a small number of outliers. Baseline drift is therefore a population phenomenon that is not dominated by a small fraction of outliers. **d** To confirm that the observed drift was not a consequence of experimental artifacts such as mechanical instability of the tissue we computed the correlation between single-cell anatomical and mean baseline ΔF/F images across trials. If drift is an anatomical artefact, excluding neurons with high positive or negative drift scores should increase correlations between anatomical images. However, distributions of correlations were similar among subsets of neurons with different drift scores. **e** Mean baseline firing rates (averaged over 20 s) of all cells (n = 7527) from all fish over trials, sorted by drift score. Firing rates are z-scored for each cell to compare each neuron’s internal trend toward more or less baseline activity over trials.

**Figure S15.**
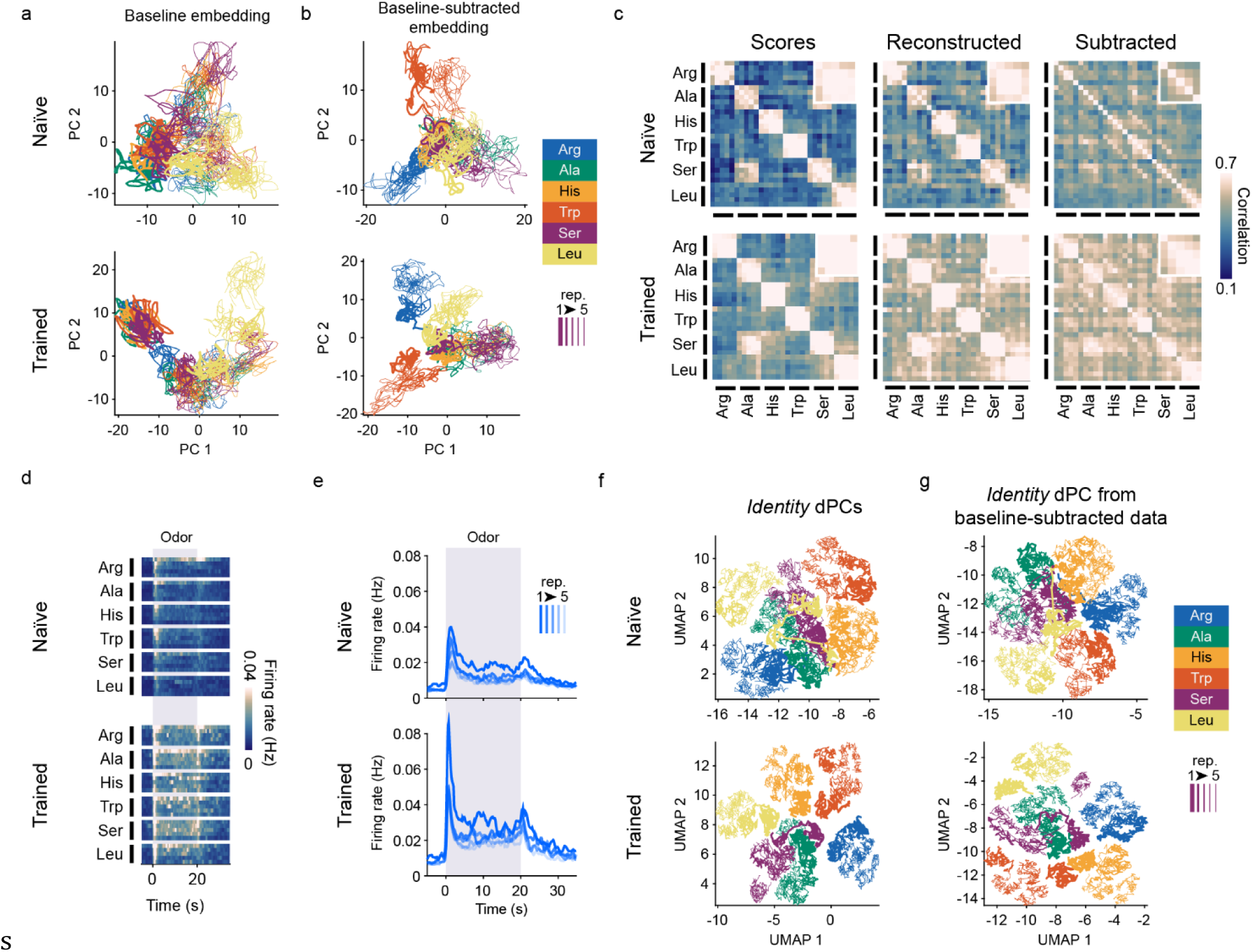
Contribution of baseline activity to representational drift, novelty and identity coding. **a** PCA of activity patterns during pre-stimulus baseline (22 to 2 seconds before onset, see also Fig. S16). Note that trajectories are organized by trial along PC1, particularly in Trained fish. **b** PCA embedding of baseline-subtracted odor responses. Note that PC1 no longer represents trials (time). Nonetheless, odor response trajectories are still clustered by odor identity and organized by repetition number within each cluster. **c-g** Results of dPCA analysis performed using baseline-subtracted data. **c** Inter-trial correlations based on identity dPCs. Left: correlations based on dPC scores. Middle: correlations based on reconstructed cellular activity. Right: correlations based on *identity*-subtracted data (raw activity – activity reconstructed from *identity* dPCs). Insets: correlations between same-odor repetitions. **d** Time course of activity (average firing rates) reconstructed from the novelty dPC only, across repetitions of each odor. Top: Naïve fish; bottom: Trained fish. **e** Average time course of reconstructed activity from novelty dPC depending on repetition number. Top: Naïve fish; bottom: Trained fish. **f-g** UMAP embedding of identity dPCs only, using full-data dPCs (**f**, see also Fig. 5d-g) or baseline-subtracted-data dPCs (**g**). Taken together, these results confirm that (1) the full-data PC1 primarily represents temporal drift, and that (2) baseline subtraction removes drift while stabilizing identity representations and preserving novelty-related components.

**Figure S16.**
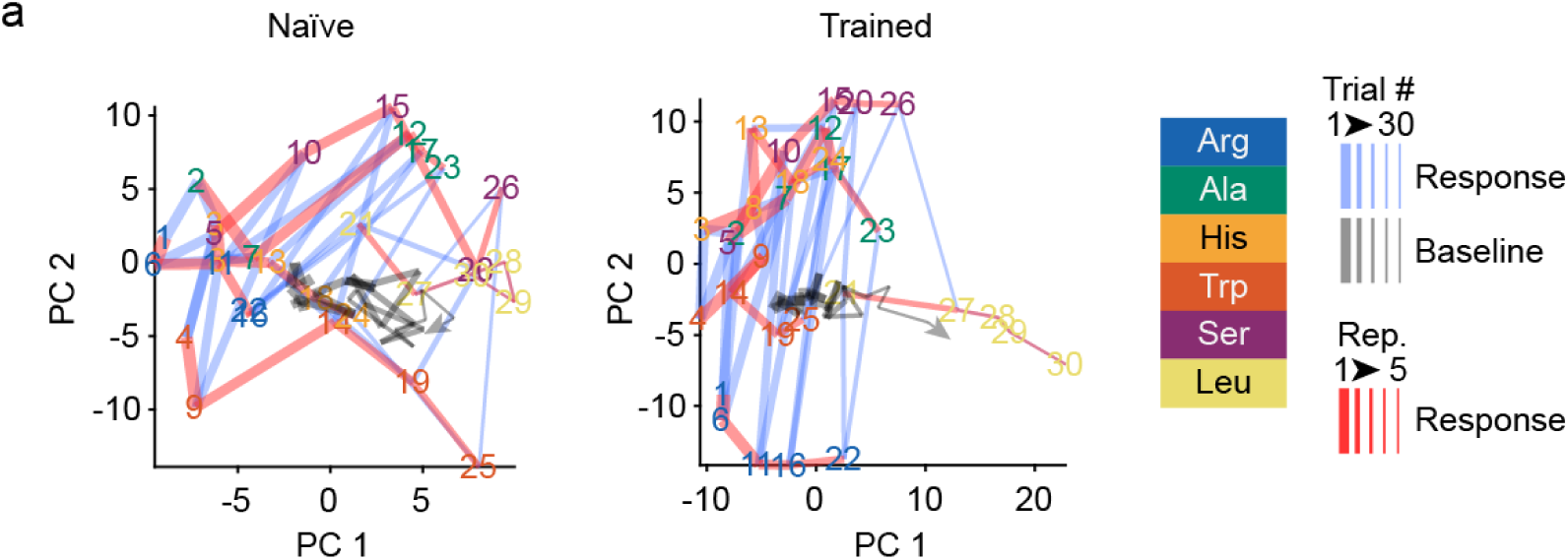
Embedding of single-trial baseline activity centroids using pre-computed PCA axes from raw odor responses. Pooled data from all Naïve (left) or Trained (right) fish. Trial numbers (colored by odor) indicate odor-response pattern centroids (same as in Fig. 7b). Each red line connects responses to consecutive repetitions of the same odor. Blue and gray lines connect successive trials (blue) or baseline patterns (gray), independent of odor identity. The gray line’s arrowhead points in the direction of time. This embedding shows the ordered arrangement of baseline patterns over trials, primarily along the PC1 axis of the odor response progression, as well as the correlation of this baseline drift axis with the average trial-to-trial drift of odor response patterns.

